# Combining genotype, phenotype, and environmental data to delineate site-adjusted provenance strategies for ecological restoration

**DOI:** 10.1101/2019.12.11.872747

**Authors:** Carolina S. Carvalho, Brenna R. Forester, Simone K. Mitre, Ronnie Alves, Vera L. Imperatriz-Fonseca, Silvio J. Ramos, Luciana C. Resende-Moreira, José O. Siqueira, Leonardo C. Trevelin, Cecilio F. Caldeira, Markus Gastauer, Rodolfo Jaffé

## Abstract

Despite the importance of climate-adjusted provenancing to mitigate the effects of environmental change, climatic considerations alone are insufficient when restoring highly degraded sites. Here we propose a comprehensive landscape genomic approach to assist the restoration of moderately disturbed and highly degraded sites. To illustrate it we employ genomic datasets comprising thousands of single nucleotide polymorphisms from two plant species suitable for the restoration of iron-rich Amazonian Savannas. We first use a subset of neutral loci to assess genetic structure and determine the genetic neighborhood size. We then identify genotype-phenotype-environment associations, map adaptive genetic variation, and predict adaptive genotypes for restoration sites. Whereas local provenances were found optimal to restore a moderately disturbed site, a mixture of genotypes seemed the most promising strategy to recover a highly degraded mining site. We discuss how our results can help define site-adjusted provenancing strategies, and argue that our methods can be more broadly applied to assist other restoration initiatives.

## Introduction

In spite of the broadly recognized importance of genetic provenance for restoration initiatives, the use of genomic tools to define provenance strategies is still uncommon. Choosing provenances based on genetic knowledge can help increase genetic diversity and adaptability, thereby contributing to the success of restoration initiatives (Broadhurst et al., 2008; Mijangos et al., 2015; Weeks et al., 2011). Fortunately, the use of genetic provenancing is increasing, as advances in Next-Generation-Sequencing technologies have made possible large-scale assessments of neutral and adaptive genetic variation (Breed et al., 2019; Mijangos et al., 2015; Williams et al., 2014). For instance, neutral loci (i.e., those which are not subject to natural selection) can be used to identify independent demographic units, assess fine-scale spatial genetic structure, and quantify genetic diversity (Allendorf et al., 2013; Balkenhol et al., 2017); whereas adaptive loci (under natural selection) are relevant to detect adaptations to local environmental conditions and delineate adaptive units (Funk et al., 2012; Rellstab et al., 2015).

Restoration genomic studies published so far have assessed the effect of multiple environmental variables on genetic composition in order to identify which individuals or populations are “pre-adapted” to future climates (Gugger et al., 2018; Lu et al., 2019; Martins et al., 2018; Rossetto et al., 2019; Shryock et al., 2017, 2015; Steane et al., 2014; Supple et al., 2018). Although this information is essential to inform predictive and climate-adjusted provenancing schemes (Prober et al., 2015), the emphasis on climate has overshadowed the application of genomic methods to restore extremely degraded sites (Bucharova et al., 2019; Lesica and Allendorf, 1999). Such site-adjusted provenancing (targeting the restoration of specific sites considering their current environmental conditions) may not even incorporate climate change considerations, as highly degraded sites have unique characteristics that make them extremely challenging to restore. First, highly degraded sites usually require immediate restoration or rehabilitation, so adaptations to current environmental conditions are more suitable to guide provenance strategies than those based on future climate (Gastauer et al., 2019). Second, environmental protection agencies usually require the restitution of ecosystems to conditions as close to a pre-disturbance baseline as possible, as well as regular (and costly) monitoring until rehabilitation goals have been achieved (Gastauer et al., 2019). The main efforts thus lay in the quick establishment of viable populations that will restore ecosystem functions and processes, prevent soil erosion, and protect biological diversity. Third, highly degraded sites such as exhausted open-pit mines have radically different environmental characteristics than natural habitats (Gastauer et al., 2019), so site-specific characteristics are of primary importance to define provenance strategies. Such site-specific variables generally need to be measured *in situ* and at fine spatial resolution, since they may not be available as spatial layers in open-access repositories (and if they are, their spatial resolution may be too coarse to reflect the reality of environmental conditions on the ground). Finally, site-adjusted provenancing strategies need to consider local adaptations to climate, soil, terrain, and even biological interactions, whereas climate-centered provenancing strategies focus exclusively on climate.

The degree of disturbance can play an important role in determining site-adjusted provenancing strategies (Breed et al., 2013; Lesica and Allendorf, 1999). Whereas local genotypes are generally the best suited to restore sites where the degree of disturbance is low, adaptations found in distant populations may facilitate establishment in highly degraded sites to which local genotypes may not be adapted (Breed et al., 2013; Broadhurst et al., 2008; Lesica and Allendorf, 1999)). Mixtures of genotypes from different populations have been suggested as the best strategy to recover highly degraded sites, given that enhanced genetic variation is more likely to rapidly generate local adaptations to novel ecological challenges (Lesica and Allendorf, 1999). In any case, determining the most appropriate site-adjusted provenancing strategy will require the delineation of local provenances and the spatial distribution of local adaptations (Breed et al., 2019).

Although the use of neutral genetic markers to identify independent demographic units is now common practice (Coates et al., 2018), few restoration studies have delineated seed sourcing strategies based on population genetic structure (Durka et al., 2017) or the genetic neighborhood size (the distance at which genetic composition stops being spatially autocorrelated) (Krauss et al., 2013; Krauss and Koch, 2004; Rossetto et al., 2019). On the other hand, only three restoration genomic studies so far have identified putative adaptive loci and then mapped adaptive genetic variation (Martins et al., 2018; Shryock et al., 2015; Steane et al., 2014). While the assessment of phenotype through common garden and reciprocal transplant experiments to identify local adaptations has a long history (Aitken and Bemmels, 2016), no study has yet combined genotype-environment associations (GEA) with genotype-phenotype associations (GPA) to delineate seed sourcing areas. This approach could improve the inference of potential candidate genes and provide important insights into genes underlying fitness-related traits (Mahony et al., 2019; Talbot et al., 2016; Vangestel et al., 2018).

Here we propose a comprehensive landscape genomic approach to assist the restoration of moderately disturbed and highly degraded sites. Relying on genotyping-by-sequencing we identified thousands of single nucleotide polymorphisms in two plant species of special interest for the restoration of exhausted mining sites from the Carajás Mineral Province, located in the Eastern Amazon (Skirycz et al., 2014; Souza-Filho et al., 2019). We first used a subset of neutral loci to assess broad and fine-scale genetic structure and determine the genetic neighborhood size. Subsequently, we combined univariate and multivariate methods to identify GEA and GPA and employed spatial principal components analyses (sPCA) to map adaptive genetic variation while accounting for spatial autocorrelation in genetic composition. Finally, we predicted the adaptive genotypes associated with the environmental conditions of restoration sites (Fig. 1).

**Figure 1.**
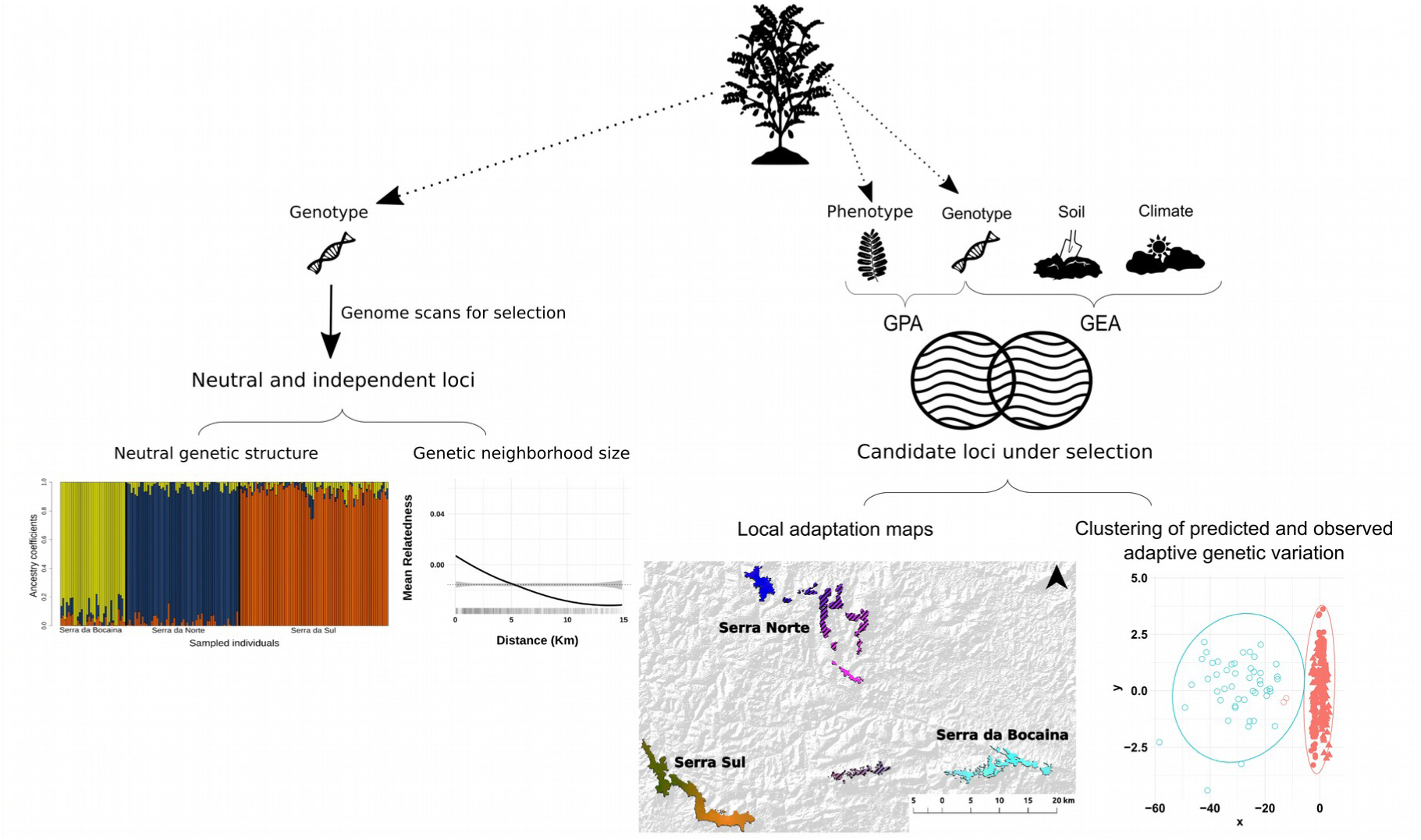
Diagram summarizing our methodological approach. We used a subset of neutral and independent loci to assess broad and fine-scale genetic structure and determine the genetic neighborhood size. Subsequently, we combined genotype, phenotype, and environmental data to identify loci under selection and then employed all candidate loci to map patterns of adaptive genetic variability and predict adaptive genotypes for restoration sites. The graphs show results for *Mimosa acutistipula.*

We focus on the Carajás Mineral Province, which harbors one of the world’s largest deposits of iron ore and huge iron ore mining projects, with operations dating back to the 1980s (Poveromo, 1999; Souza-Filho et al., 2019). The banded ironstone formations known as *Cangas* (where iron-ore deposits are concentrated) are characterized by shallow, acidic, nutrient-depleted and metal-rich soils and marked by high solar radiation, hot temperatures, and a severe drought period (Skirycz et al., 2014), which pose severe challenges to plant growth. Environmental legislation in Brazil requires the rehabilitation of disabled iron ore mining sites (Gastauer et al., 2019), which constitute extremely degraded and difficult to restore environments. These mining sites are characterized by extensive vegetation and soil removal, compacted, nutrient-poor soils and steep slopes (Boyer and Wratten, 2010; Garris et al., 2016; Whiting et al., 2004). The successful restoration and rehabilitation of these highly degraded sites thus requires an appropriate selection of plant species (Giannini et al., 2017) and seed sourcing zones to ensure that the introduced plants can effectively colonize and establish viable populations. However, mine rehabilitation programs in the region employ seed mixtures of exotic plants, due to the scarcity of native seeds and their lower germination and growth rates (Silva et al., 2018). Additionally, some natural Canga environments from the region have been repeatedly disturbed by fires, illegal cattle ranching and the introduction of alien plant species, such as grasses and ferns. This is the case in Serra da Bocaina, which became a National Park in 2017 and has since been protected (Mota et al., 2018). No restoration initiatives have yet been implemented to recover the original ecosystems found in Serra da Bocaina, but provenance strategies are expected to differ substantially from those required to rehabilitate disabled mines.

Being native species, dominant in Canga environments, and once found at the sites being restored, *Mimosa acutistipula var. ferrea* Barneby and *Dioclea apurensis* Kunth (both legumes) are among the most promising plants for use in Canga restoration and mineland rehabilitation programs (Giannini et al., 2017). Considered metallophyte species, both exhibit biological mechanisms to tolerate and thrive in metalliferous soils (Preite et al., 2019; Whiting et al., 2004). Moreover, they are extremely abundant in pristine Cangas ecosystems and interact symbiotically with nitrogen-fixing bacteria, thus contributing to soil enrichment and acting as pioneer species in restoration sites (Nunes et al., 2015; Ramos et al., 2019a; Silva et al., 2018). *Mimosa acutistipula* is drought tolerant and well adapted to the low nutrient content of Canga soils (Silva et al., 2018). On the other hand, *D. apurensis* requires low nutrient inputs and shows high nutrient use efficiency (Ramos et al., 2019b). Moreover, this species is a fast-growing liana with a ground-covering growth form, enabling the revegetation and stabilization of mine pits and waste piles. Both species have high germination rates (Ramos et al., 2019a), can be observed growing on minelands, and seem to be central in plant-pollinator networks (unpublished data). Considering the heterogeneous and hostile environment where both species occur (Mitre et al., 2018), and their life-history similarities, we expected to find similar patterns of neutral and adaptive genetic structure in both species across our study area. We also expected that local populations would not be adapted to the environmental conditions found in exhausted mining sites given they drastically differ from pre-mining conditions, whereas populations from Serra da Bocaina would show adaptations to local environmental conditions. Based on our results, we propose two different site-adjusted provenance strategies for the restoration of a degraded mine site and a disturbed but unmined Canga environment. We discuss the merits of our approach and argue that it can be more broadly applied to define site-adjusted provenancing strategies.

## Material and Methods

### Sampling

We followed a stratified sampling design, seeking to ensure high statistical power in GEA and GPA analyses by maximizing environmental variability within different genetic clusters. We collected samples of 180 individuals of *M. acutistipula* var. *ferrea* and 167 individuals of *D. apurensis* between February and May of 2018 (SISBIO collection permit N. 48272-6), across the three major Canga highlands of the Carajás Mineral Province (Fig. 2). For each individual plant, we collected a sample of root-proximal soil (0-5 cm) for chemical characterization and leaflet samples for phenotype and genotype analyses. These Canga ecosystems are composed of several physiognomies, comprising grasslands, scrublands, wetlands and forest formations (Mota et al., 2015), which differ in terms of the plant communities they support as well as in their soil chemistry (Mitre et al., 2018). To ensure sampling across environmental gradients, individuals were collected in each one of these physiognomies within each highland. We also scattered samples to cover the full extent of each highland (Fig. 2). A minimum distance of 20 meters between samples was used to minimize sampling related individuals. In addition to the soil samples gathered along with plant tissue, we collected 50 extra soil samples from a highly degraded mine site (the mine pit from an exhausted mine) and seven soil samples from a never mined, but moderately disturbed site (scattered across Serra da Bocaina, Fig. 2). These soil samples were used to predict adaptive genotypes associated with the environmental conditions of both restoration sites (see details below).

**Figure 2.**
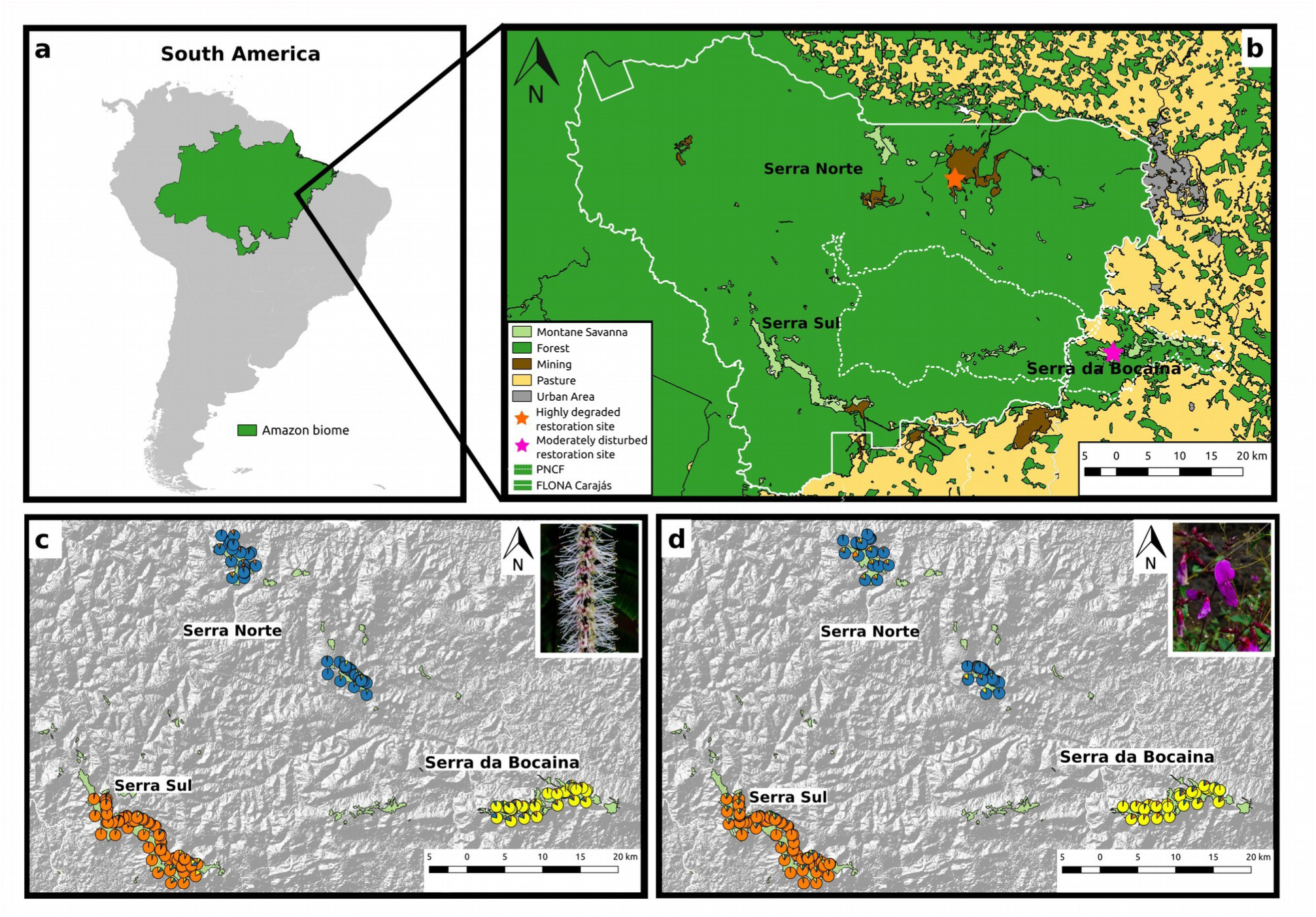
Maps of our study region depicting restoration sites and neutral genetic structure for *Mimosa acutistipula* var. *ferrea* and *Dioclea apurensis.* **a**: Location of the Carajás Mineral Province within the Amazon biome. **b:** Location of the major Canga highlands (Montane Savanna) where samples were collected. White continuous lines represent the Carajás National Forest (FLONA Carajás), white dashed lines the Campos Ferruginosos National Park (PNCF), and stars depict restoration sites. Ancestry coefficients for all samples from *M. acutistipula* var. *ferrea* (**c**) and *D. apurensis* (**d**), determined using the *snmf* function from the LEA package.

### Environmental data

Soil samples were air-dried and sieved using 2 mm mesh, and once dry they were sent to LABRAS (http://labrasambientaiseagricolas.com.br/) for chemical analyses. These included pH, organic matter, available P, K, and Na, exchangeable Ca, Mg, and Al, exchangeable S and available B, Cu, Fe, Mn and Zn (see details in Supporting Information Methods S1). To reduce the number of chemical parameters describing soil composition, we selected those known to most affect plant physiology in metalliferous ecosystems (organic matter, Fe, Mn, P, pH, S, and B (Bothe, 2011; Mitre et al., 2018; Whiting et al., 2004)), along with a set of orthogonal variables explaining most variation in soil composition across our study area. To identify this set of orthogonal variables we first used the function *imputePCA* from the *missMDA* R package (Josse and Husson, 2016) to impute missing data (20 samples contained missing data for at least one parameter) using the regularized iterative PCA algorithm recommended to avoid overfitting (Josse and Husson, 2016). We then ran separate principal component analyses (PCA) for each species using all the centered and scaled chemical parameters and selected the three variables showing the strongest correlation with the first, second and third PC axes (each showing eigenvalues > 1 and > 10% of total variance explained, and all three explaining > 50% of total variance; Table S1). The selected soil variables were organic matter, Zn and Na in both species).

We also retrieved climatic data from WorldClim version 1 [1950-2000; (Hijmans et al., 2005)], using the sample coordinates to extract all bioclimatic variables. We followed a similar protocol to obtain the set of orthogonal variables explaining most climatic variance across our study region (Table S1). The first three PCA axes explained 80% of total climatic variance in both species, and the bioclimatic variables most strongly correlated with those axes were isothermality (bio03), minimum temperature of coldest month (bio06) and precipitation of driest quarter (bio17) for *M. acutistipula*; and isothermality (bio03), minimum temperature of coldest month (bio06) and maximum temperature of warmest month (bio05) for *D. apurensis* (see Fig. S1 for maps of these layers). Correlations between these environmental variables were all below |r| < 0.6 (Fig. S2).

### Phenotypic data

For each leaf sample we determined macro- and micro-nutrient content and the specific leaf area (SLA), using standard methods (see details in Methods S1). We selected those phenotypic variables known to affect plant physiology in metalliferous ecosystems (SLA, N, B, Fe, Mn, P, N/P, (Bothe, 2011; Mitre et al., 2018; Pérez-Harguindeguy et al., 2013). As described above, we also selected the three orthogonal phenotypic variables explaining most of phenotypic variance (the first three PCA axes explained more than 50% of total variance in both species, Table S1) after imputing missing data (11 samples contained missing data in at least one parameter). Selected phenotypic variables were Zn, N and B for *M. acutistipula* and P, Mn and K for *D. apurensis* (correlations between these phenotypic variables were all below |r| < 0.6; Fig. S2).

### Genome size estimation, DNA extraction, genotype-by-sequencing and bioinformatic processing

We used flow cytometry to estimate haploid genome size in both species (1C DNA content was 712 Mbp in *M. acutistipula* and 642 Mbp in *D. apurensis*, see details in Methods S1). Total DNA was extracted using Qiagen’s DNeasy Plant Mini Kit. DNA concentration was quantified using Qubit High Sensitivity Assay kit (Invitrogen), and DNA integrity assessed through 1.2% agarose gel electrophoresis. Samples with concentrations below 5 ng/µL or showing no clean bands were excluded from all analyses, and selected samples were normalized to a concentration of 5 ng/µL and a total volume of 30 µL. These were then shipped to SNPSaurus (http://snpsaurus.com/) for sequencing and raw data bioinformatic processing (see details in Methods S1). Briefly, genomic DNA was converted into nextRAD genotyping-by-sequencing libraries (SNPsaurus, LLC) as in (Russello et al., 2015), considering the estimated genome size of each species. Genomic DNA was first fragmented with the Nextera DNA Flex reagent (Illumina, Inc), which also ligates short adapter sequences to the ends of the fragments. The Nextera reaction was scaled for fragmenting 14 ng and 20 ng of genomic DNA for *M. acutistipula* and *D. apurensis*, respectively. Fragmented DNA was then amplified for 25 cycles at 75 degrees, with one of the primers matching the adapter and extending 8 nucleotides into the genomic DNA with the selective sequence TGCAGGAG. Thus, only fragments starting with a sequence that can be hybridized by the selective sequence of the primer were efficiently amplified. The nextRAD libraries were then sequenced on a HiSeq 4000 with six and five lanes of 150 bp reads for *M. acutistipula* and *D. apurensis*, respectively (University of Oregon). Reads were trimmed using *BBMap tools* (http://sourceforge.net/projects/bbmap/) to exclude Nextera adapters and a reference contig was created by collecting 10 million reads in total, evenly distributed from the samples, and excluding reads that had counts fewer than 6 or more than 800. The remaining loci were then aligned to each other to identify allele loci and collapse allelic haplotypes to a single representative. All reads were mapped to the reference contig with an alignment identity threshold of 95% using *BBMap tools*. Genotype calling was done using *callvariants* (*BBMap tools*), and the resulting set of genotypes were filtered to remove alleles with population frequency of less than 3%. Loci that were heterozygous in all samples and loci that contained more than 2 alleles in a sample (suggesting collapsed paralogs) were removed. A total of 7,165 RAD-tag sequences were obtained for *M. acutistipula* and 4,325 for *D. apurensis.* Considering the genome size of each species and a linkage block size of 378 Kbp (mean value for the Fabaceae family, (Lowry et al., 2017)), we estimated a maximum proportion of genome coverage (assuming one RAD-tag per block) of 100% (McKinney et al., 2017). From those RAD-tags, 17,403 SNPs were generated for *M. acutistipula* and 9,857 SNPs for *D. apurensis* (minimum sequencing depth of 14 and 9, respectively).

### Genetic diversity and neutral genetic structure

The R package *r2vcftools* (https://github.com/nspope/r2vcftools) - a wrapper for VCFtools (Danecek et al., 2011) - was used to perform final quality control on the genotype data. To assess neutral genetic structure and genetic diversity, we used a series of filters to obtain a set of neutral and independent loci. Filtering criteria included quality (Phred score > 30), read depth (20 – 800), minor allele frequency (MAF > 0.05), linkage disequilibrium (*r*2 < 0.8, (Xuereb et al., 2018)), Hardy-Weinberg Equilibrium (HWE, *p* > 0.0001), and loci and individuals with less than 20% missing data (an example filtering script can be seen in https://github.com/rojaff/r2vcftools_basics). Additionally, we removed loci potentially under selection using genome scans. These accounted for population structure (assessed using the *snmf* function from the *LEA* package, as described below), and controlled for false discovery rates by adjusting *p-*values with the genomic inflation factor (λ) and setting false discovery) and setting false discovery rates to *q*=0.05, using the Benjamini-Hochberg algorithm (François et al., 2016) (see details below).

We used two complementary genetic clustering approaches to assess neutral population structure: the *snmf* function from the *LEA* package (Frichot and François, 2015), and Discriminant Analysis of Principal Components - DAPC from the *adegenet* package (Jombart and Ahmed, 2011). The *snmf* model implements a fast yet accurate likelihood algorithm (Frichot et al., 2014), while DAPC is a robust genetic clustering method with no assumption about the underlying population genetic model (Jombart and Ahmed, 2011). Based on previous population genomic studies for other co-occurring plant species (Carvalho et al., 2019; Lanes et al., 2018; Silva et al., 2020), we tested from one to ten ancestral populations (*k*). In the case of *snmf* we performed ten replicate runs for each value of *k*, choosing the most likely *k* based on minimized cross-entropy. For DAPC, we inferred optimal *k* using k-means clustering and the Bayesian Information Criterion (BIC). Considering the ancestry coefficients assigned by *snmf*, we then estimated expected heterozygosity (*HE*), inbreeding coefficients (*F*), and nucleotide diversity (π) for each genetic cluster. We also estimated pairwise *FST* using dartR R package (Gruber et al., 2018), and effective population sizes (*Ne*) employing the linkage disequilibrium method implemented in NeEstimator 2.1 and a lowest allele frequency value of 0.05 (Do et al., 2014). Finally, we assessed fine-scale spatial genetic structure in each species within each genetic cluster through local polynomial fitting (LOESS) of Yang’s genetic relatedness between pairs of individuals (Yang et al., 2010) and pairwise geographic distance, as in (Carvalho et al., 2019).

### Assessing genotype-environment associations (GEA) and genotype-phenotype associations (GPA)

To assess GEA and GPA (Fig. 1) we first filtered loci for quality (Phred score > 30), read depth (20-800), minor allele frequency (MAF > 0.05), linkage disequilibrium (*r²* < 0.8), and loci and individuals with less than 20% of missing data. We then combined univariate and multivariate methods, namely Latent Factor Mixed Models (LFMM) and Redundancy Analysis (RDA). While LFMM identifies associations between single loci and single predictors, RDA can detect multilocus signatures of selection as a function of a multivariate set of predictors (Caye et al., 2019; Forester et al., 2018). Both methods assume a linear relationship between allele frequency and environmental variables, have been used extensively (Ahrens et al., 2018), provide a good compromise between detection power and error rates, and are robust to a variety of sampling designs and underlying demographic models (Forester et al., 2018; Rellstab et al., 2015). Since both methods require complete data sets (without missing values), we performed an imputation of missing genotypes (7.6% and 7% missing genotypes for *M. acutistipula* and *D. apurensis* respectively) based on the *snmf* population assignments from the previous step, using the *impute* function and the mode method from the *LEA* package (Frichot and François, 2015). This function imputes missing genotypes using ancestry and genotype frequency estimates from the *snmf* run.

LFMM analysis were performed using the *lfmm* package (Caye et al., 2019) and ridge estimates, which minimize regularized least-squares with a *L2* penalty (see example script here: https://bcm-uga.github.io/lfmm/articles/lfmm). Instead of using raw predictor variables, we employed the first four axes resulting from a Principal Components Analysis (PCA) on all predictor variables in order to minimize the number of tests. These four axes explained more than 60% of total environmental and phenotypic variance in both species, and were strongly correlated (|r| > 0.7) with organic material, B, Fe, Bio06, Bio17, Zn, S and Na (environmental variables), and N/P, P, Fe (phenotypic variables) in both species. We ran LFMM using the previously identified number of genetic clusters (*k*=3, see results) as latent factors, to account for the underlying neutral genetic structure. We then calculated the genomic inflation factor (λ) and setting false discovery) and modified it until a calibrated distribution of adjusted *p-*values was found, and set false discovery rates at a rate of *q*=0.05 using the Benjamini–Hochberg algorithm (François et al., 2016).

We performed RDA using the *rda* function from the *vegan* package (Oksanen et al., 2019) as implemented in Forester et al. (2018), modeling genotypes as a function of predictor variables, and producing as many constrained axes as predictors (see example script here: https://popgen.nescent.org/2018-03-27_RDA_GEA.html). Multicollinearity between predictors was assessed using the variance inflation factor (VIF) and since all predictor variables showed VIF < 3 none were excluded. Raw predictor variables were scaled and centered prior to analyses and the population assignments from *snmf* (population ID) were used to control for population structure by running a partial RDA. Significance of RDA constrained axes was assessed using the *anova.cca* function and significant axes were then used to identify candidate loci in both species. Candidate loci were identified using a Mahalanobis distance-based approach (Capblancq et al., 2018), which made RDA result comparable with those obtained with LFMM, since it allowed adjusting *p-*values using the genomic inflation factor (λ) and setting false discovery) and setting false discovery rates to *q*=0.05, as described above (calculated and modified genomic inflation factors and *p*-value distributions for LFMM and RDA tests are provided in Figs. S3-S8). To assess the impact of population genetic structure on our number of detections we ran additional cluster-level GEA analyses (LFMM and RDA), using only individuals belonging to the same genetic cluster (setting *k*=1 in LFMM and omitting population ID in RDA). Finally, to visualize patterns of GEA and GPA, we ran additional RDA models excluding neutral loci, using the combined candidate adaptive loci detected using the general RDA and LFMM analyses.

In order to search for the proteins coded by the genes contained in the flanking regions of our candidate SNPs (found in GEA and GPA analyses), contig sequences containing candidate loci were first submitted to the EMBOSS Transeq (http://www.ebi.ac.uk/Tools/st/emboss_transeq/) to obtain corresponding protein sequences. We used all six frames with standard code (codon table), regions (start-end), trimming (yes), and reverse (no). We then ran a functional analysis using InterPro (https://www.ebi.ac.uk/interpro/; interproscan.sh -dp –appl PfamA, TIGRFAM, PRINTS, PrositePatterns, Gene3d –goterms –pathways -f tsv -o MySequences.tsv -i MySequences.faa), searching for gene ontology terms and pathways along a variety of annotation databases (i.e., Interpro, Pfam, Tigrfam, Prints, PrositePattern and Gene3d).

### Mapping adaptive genetic variation

To map adaptive genetic variation, we used the *adegenet* package (Jombart and Ahmed, 2011) to run a Spatial Principal Component Analysis (sPCA) on the combined candidate adaptive loci detected in GEA and GPA analyses using general LFMM and RDA (results for intersected loci are presented in Fig. S15). sPCA is a spatially explicit multivariate method that yields scores summarizing genetic variability and spatial structure among individuals (Jombart et al., 2008). Spatial structure is estimated using a Moran’s Index that relies on the comparison of allelic frequencies observed in one individual to the values observed in neighboring individuals. These neighboring individuals can be defined by distinct connection networks, which in our case was set to a distance-based neighborhood, as indicated for aggregated distributions (Jombart et al., 2008). The Moran’s Index generates two types of spatial structuring: global structure, which reflects positive spatial autocorrelation, and local structure, that reflects negative spatial autocorrelation (Jombart et al., 2008). To decide if global and/or local structures should be interpreted and thus retained in sPCA analyses, we used the global and local tests proposed by Jombart & Ahmed (2011). The first three retained axes were then interpolated on 10 meter resolution grids covering our study area, and the resulting rasters used to create an RGB composite, using the Merge function in QGIS 3.4 (see example scripts here: https://github.com/rojaff/LanGen_pipeline). The resulting color patterns represent the similarity in adaptive genetic composition.

To predict the adaptive genotypes associated with environmental data collected from restoration sites (the highly degraded exhausted mine and the moderately disturbed Serra da Bocaina, Fig. 2), we employed the GEA-RDA models fitted on the combined candidate adaptive loci detected by global LFMM and RDA (see previous section), and ran the *predict.cca* function from the *vegan* package. Environmental samples (soil and climate) from these sites were thus used to predict RDA scores, based on the fitted GEA-RDA models. We then performed a *k*-means clustering analysis (using Euclidean distances) on observed and predicted RDA scores for individuals from each species, using all significant constrained axes and allowing the number of clusters to vary between two and five (three Canga highlands and two restoration sites). We used the *NbClust* package (Charrad et al., 2014) to obtain the optimal number of clusters chosen by 30 different algorithms. Observed and predicted RDA scores groupping together, suggest that our sampled individuals possess adaptations associated with the environmental conditions of restoration sites. Observed and predicted RDA scores placed in different clusters, on the other hand, indicate that none of our sampled individuals seems adapted to the environmental conditions of restoration sites.

## Results

### Genetic diversity and neutral genetic structure

After filtering for quality, read depth, minor allele frequencies, missing data, linkage disequilibrium, Hardy-Weinberg Equilibrium, and outlier loci, we retained 7,376 and 3,496 neutral and independent SNPs and 177 and 163 individuals for *M. acutistipula* and *D. apurensis*, respectively, which were then used to assess genetic diversity and population structure. Both genetic clustering approaches (*snmf* and DAPC) indicated the presence of three clusters in the two study species (Fig. S9). Admixture levels were low, all individuals were correctly assigned to their source Canga highland (Fig. 2), and there was genetic differentiation between genetic clusters (pairwise *FST* values were significant and ranged between 0.11 and 0.13 in *M. acutistipula* and between 0.16 and 0.27 in *D. apurensis*). Expected heterozygosity and nucleotide diversity were similar in both species, but inbreeding coefficients were lower and effective population sizes larger in *M. acutistipula* (Table 1). Both species showed significant inbreeding coefficients in all genetic clusters and exhibited the largest effective population sizes in Serra Sul (Table 1). We detected spatial autocorrelation in genetic relatedness within genetic clusters in each species (Fig. S10-S11). In both, the strength of spatial autocorrelation was highest in Serra Sul, where genetic neighborhood size was larger (∼5km, Fig. S10-S11).

**Table 1.**
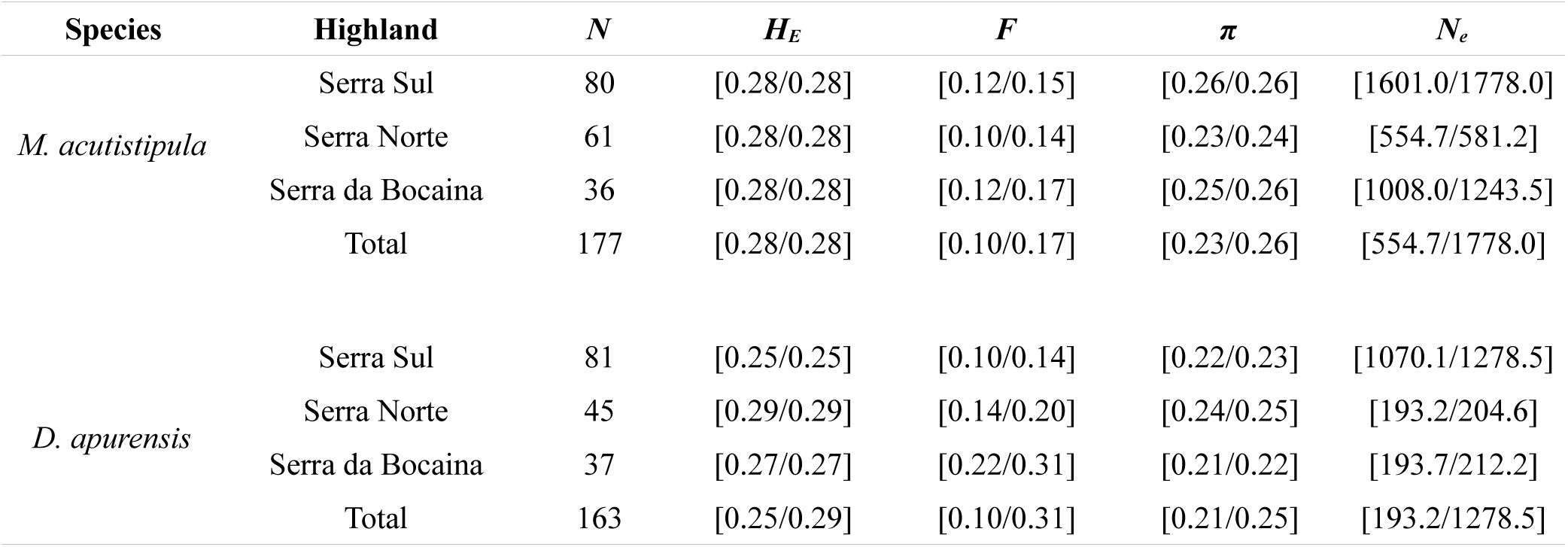
Genetic diversity measures for *Mimosa acutistipula* var. *ferrea* and *Dioclea apurensis* within genetic clusters (highlands). Sample sizes (*N*) are followed by mean expected heterozygosity (*HE*), mean inbreeding coefficient (*F*), nucleotide diversity (π) and effective population size (*Ne*). Values represent 95% confidence intervals.

| Species | Highland | $N$ | $H_E$ | $F$ | $\pi$ | $N_e$ |
| --- | --- | --- | --- | --- | --- | --- |
| *M. acutistipula* | Serra Sul | 80 | [0.28/0.28] | [0.12/0.15] | [0.26/0.26] | [1601.0/1778.0] |
|  | Serra Norte | 61 | [0.28/0.28] | [0.10/0.14] | [0.23/0.24] | [554.7/581.2] |
|  | Serra da Bocaina | 36 | [0.28/0.28] | [0.12/0.17] | [0.25/0.26] | [1008.0/1243.5] |
|  | Total | 177 | [0.28/0.28] | [0.10/0.17] | [0.23/0.26] | [554.7/1778.0] |
| *D. apurensis* | Serra Sul | 81 | [0.25/0.25] | [0.10/0.14] | [0.22/0.23] | [1070.1/1278.5] |
|  | Serra Norte | 45 | [0.29/0.29] | [0.14/0.20] | [0.24/0.25] | [193.2/204.6] |
|  | Serra da Bocaina | 37 | [0.27/0.27] | [0.22/0.31] | [0.21/0.22] | [193.7/212.2] |
|  | Total | 163 | [0.25/0.29] | [0.10/0.31] | [0.21/0.25] | [193.2/1278.5] |

### Genotype-environment and genotype-phenotype associations

After filtering for quality, read depth, minor allele frequencies, missing data, and linkage disequilibrium we retained 9,480 and 4,720 SNPs and 177 and 163 individuals for *M. acutistipula* and *D. apurensis*, respectively. Using LFMM we identified a total of 198 and 154 contigs (RAD-tags) containing GEA, and 94 and 185 contigs containing GPA in *M. acutistipula* and *D. apurensis*, respectively (Tables S2 and S3). Only the first two constrained axes from RDA analyses were significant (ANOVA’s *p* < 0.05) in GEA and GPA analyses for both species. RDA revealed a total of 403 and 225 contigs containing significant GEA and 281 and 119 contigs containing significant GPA in *M. acutistipula* and *D. apurensis* respectively (Fig. 3, Fig. S12 and Tables S2 and S3). In *M. acutistipula* 344 contigs were most correlated to climatic variables and 69 to soil variables, while in *D. apurensis* 203 contings were most correlated to climatic and 23 to soil variables. Combining both methods (LFMM and RDA), we found a total of 588 contigs showing GEA in *M. acutistipula* and 360 in *D. apurensis*, and 368 contigs showing GPA in *M. acutistipula* and 288 in *D. apurensis*. Only 108 contigs contained both GEA and GPA in *M. acutistipula* and 65 in *D. apurensis.* Finally, cluster-level GEA analyses revealed many cluster-exclusive detections in both species (Fig. S13).

**Figure 3.**
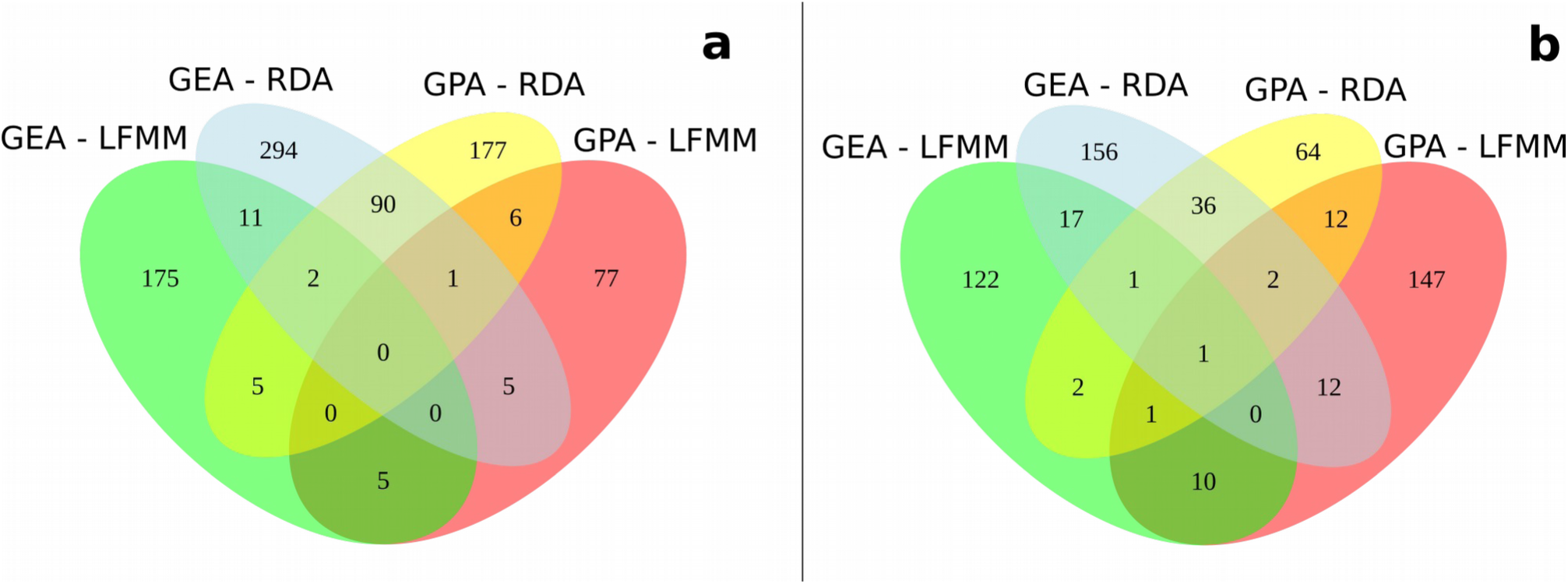
Venn diagram showing the intersection of contigs (RAD-tags) containing candidate SNPs for *Mimosa acutistipula* var. *ferrea* (**a**) and *Dioclea apurensis* (**b**). Genotype-environment associations (GEA) and genotype-phenotype association (GPA) were assessed using Redundant Analysis (RDA) and Latent Factor Mixed Model (LFMM). The number of detections by method for each species are presented in Tables S2 and S3.

Subsequent RDA models using the combined candidate adaptive loci detected using general LFMM and RDA analyses revealed population-level patterns of GEA and GPA (Fig. 4). In *M. acutistipula* GEA and GPA models explained 17% and 5% of total variance respectively, while in *D. apurensis* GEA and GPA models explained 31% and 9% of total variance. In both species, axes loadings were higher for climatic variables (0.01-0.89 for *M. acutistipula* and 0.01-0.83 for *D. apurensis*) than for soil variables (0.005-0.47 for *M. acutistipula* and 0.003-0.59 for *D. apurensis*). In *M. acutistipula*, the first and second axes split individuals into three large GEA groups corresponding to their sampling location. While individuals from Serra Norte showed associations with higher isothermality (bio03) and higher winter temperatures (bio06), individuals from Serra Sul showed associations with warmer winter temperatures and wetter dry season precipitation (bio17). Individuals from Serra da Bocaina exhibited associations with higher pH (less acidic soils) and drier dry season precipitation (Fig. 4a). Interestingly, *Dioclea apurensis* showed similar GEA patterns based on isothermality (bio03), winter temperatures (bio06), and pH, despite using a slightly different set of predictors (Fig. 4b). On the other hand, the first and second constrained axes divided individuals into two large GPA groups in *M. acutistipula* (Fig. 4c), the first one encompassing individuals from Serra Norte (which showed associations with higher SLA and Mn, and lower P), and the second individuals from Serra Sul and Serra da Bocaina (showing associations with lower SLA and Mn). In *D. apurensis*, the first and second axes split individuals into three GPA groups, with individuals from Serra Norte showing associations with a higher leaf content of Fe and Mn and a lower content of P, while those from Serra Sul showed associations with higher N and those from Serra da Bocaina with lower SLA and N/P (Fig. 4d). Leaf-level nutrients were weakly correlated with soil-level nutrients (Pearson’s correlation coefficients ranged between −0.07 and 0.39 for *M. acutistipula* and between −0.04 to 0.24 for *D. apurensis*).

**Figure 4.**
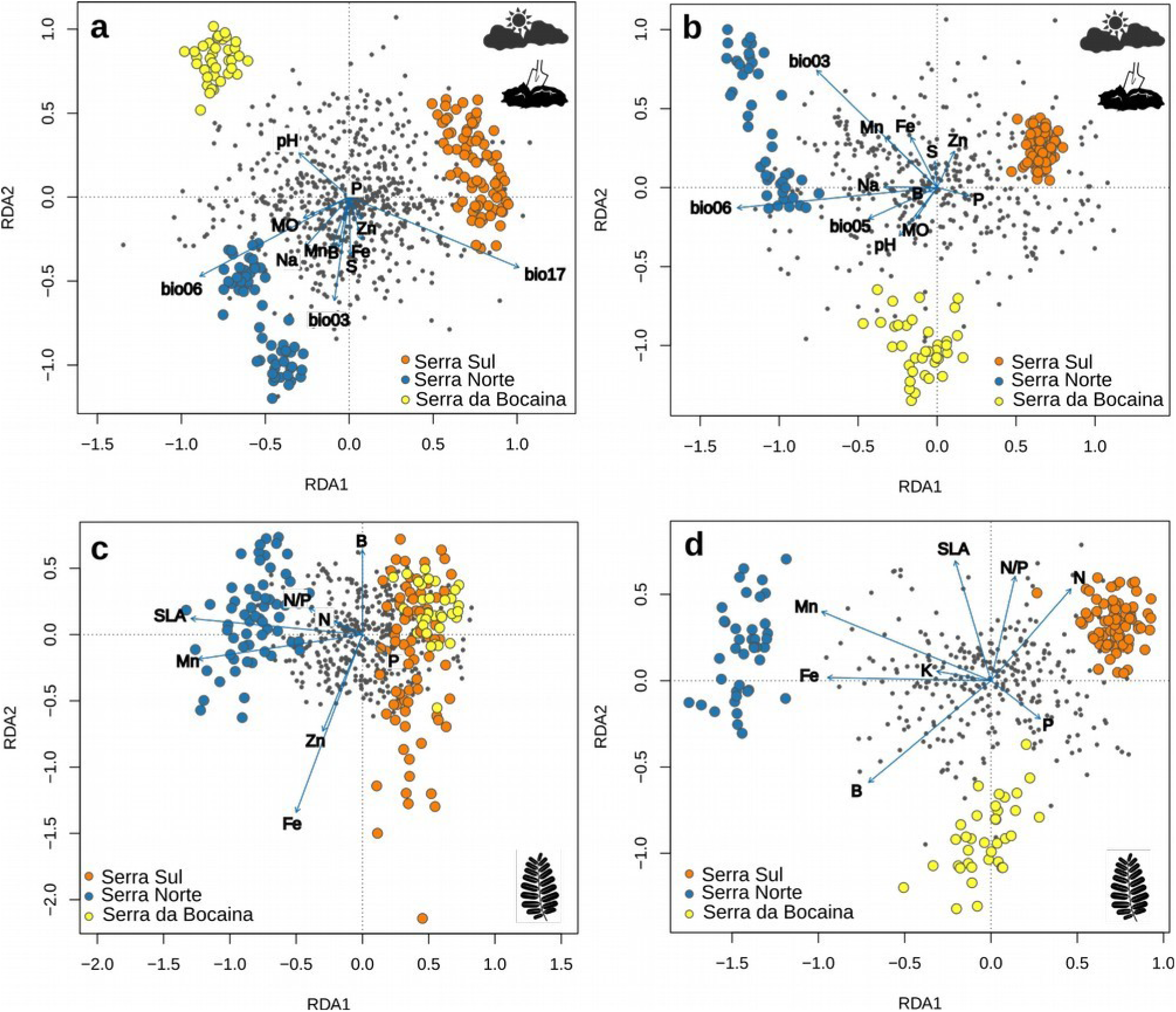
Redundancy analysis (RDA) using only the candidate loci identified in genotype-environment (upper panels) and genotype-phenotype association analyses (lower panels) in *Mimosa acutistipula* var. *ferrea* (**a**, **c**) and *Dioclea apurensis* (**b**, **d**). The plots show the first and second constrained axes from RDA, with SNPs represented as gray filled circles, environment and phenotype variables as blue arrows, and individuals from different Canga highlands in color circles (bio03: isothermality; bio05: maximum temperature of warmest month; bio06: minimum temperature of coldest month; bio17: precipitation of driest quarter; MO: organic material; and SLA: specific leaf area).

A subset of the contigs containing candidate SNPs showed InterPro annotations (105 contigs in *M. acutistipula* and 59 in *D. apurensis*). Candidate adaptive genes were associated to different functions, including intracellular transport, catalytic activity, synthesis of hormones, metabolic and oxidation-reduction processes, and plant defense response (a full list of candidate genes with InterPro annotations is presented in Table S4). Only 17 putative adaptive genes containing InterPro annotations were shared between both species (Table S5).

### Mapping adaptive genetic variation

The combination (union) of candidate adaptive loci detected through GEA and GPA resulted in 914 loci for *M. acutistipula* and 614 loci for *D. apurensis.* Since none of the sPCA local structure tests were significant, we retained the first three positive global axes, which explained most variance in both species (51% and 81% of total variance for *M. acutistipula* and *D. apurensis* respectively, Fig. S14). These revealed a similar adaptive genetic structure in both species (Fig. 5), with two adaptive units in Serra Norte and one in Serra da Bocaina. *Mimosa acutistipula* nevertheless exhibited a clinal adaptive pattern in Serra Sul, whereas *D. apurensis* did not. Similar spatial patterns were found when using the intersected loci (i.e. those shared by GEA and GPA; Fig. S15). Finally, predicted genotypes associated with climatic and soil characteristics from a highly degraded mining site did not cluster together with any of our study populations in either species (Fig. 6a and 6b). In contrast, most predicted genotypes for the environmental conditions from the moderately disturbed Serra da Bocaina clustered together with individuals collected in the same location, revealing local provenances are putatively adapted to local environmental conditions (Fig. 6c and 6d).

**Figure 5.**
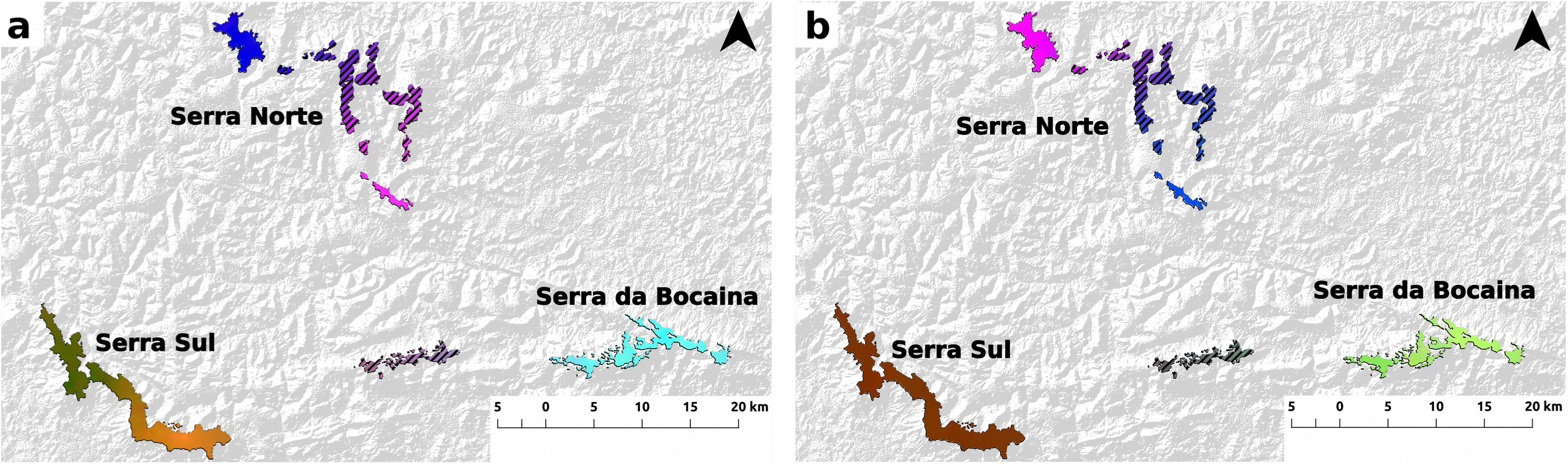
Spatial distribution of adaptive genetic variation in *Mimosa acutistipula* var. *ferrea* (**a**) and *Dioclea apurensis* (**b**). The maps represent an RGB composite made using interpolated principal components from a sPCA, ran on the combined candidate loci found in GEA and GPA. Regions with similar colors within each panel represent analogous genetic composition and areas with diagonal lines were not sampled (i.e., adaptive genetic composition was extrapolated from neighboring areas).

**Figure 6.**
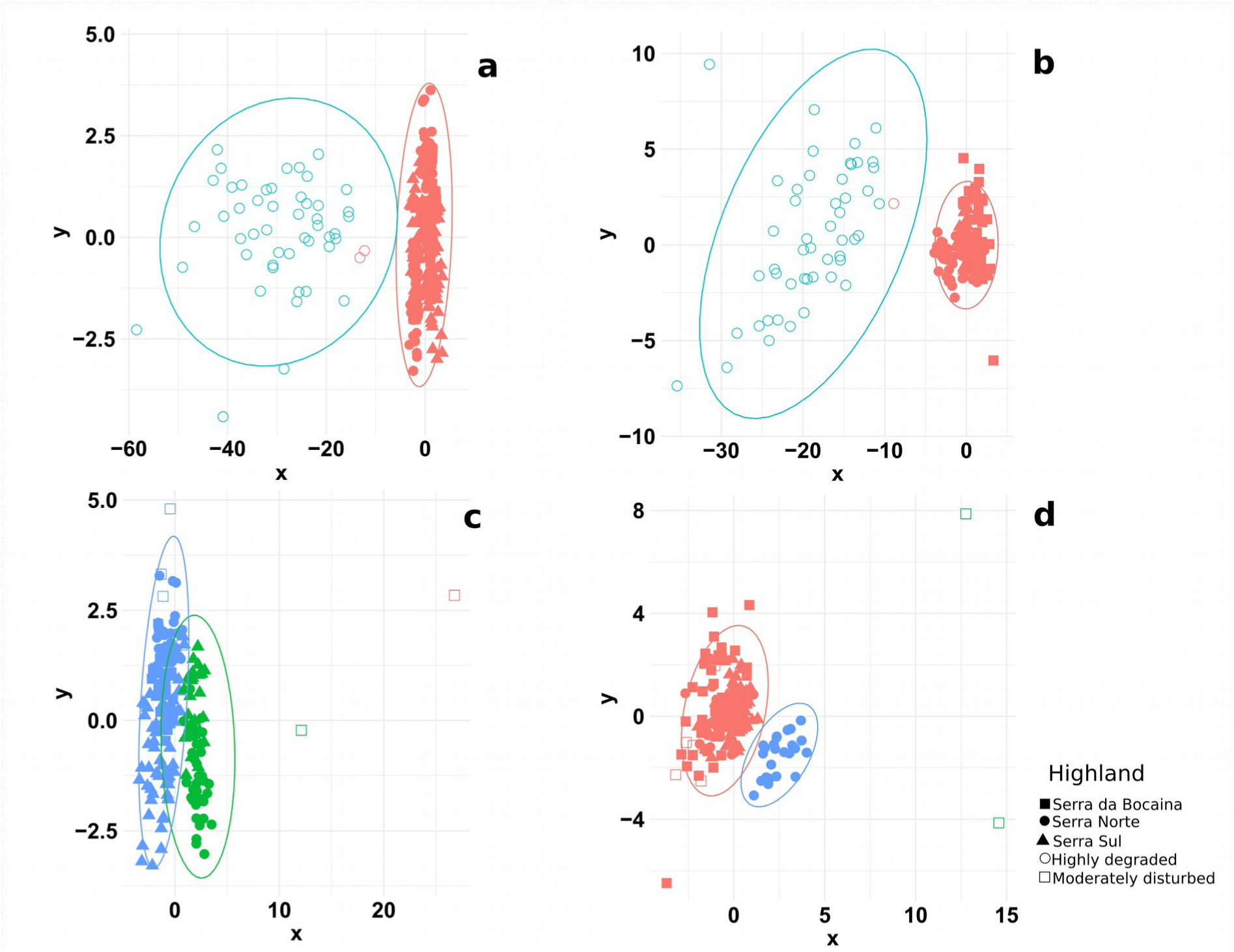
Clustering of predicted and observed adaptive genetic variation in *Mimosa acutistipula* var. *ferrea* (a, c) and *Dioclea apurensis* (b, d). A *k*-means clustering approach was employed to examine the similarity between observed (filled symbols) and predicted genotypes (open symbols) associated with the environmental conditions of highly degraded (upper panels) and moderately disturbed (lower panels) sites. Colors indicate different clusters and symbols the locations where samples were collected.

## Discussion

The delineation of seed sourcing areas requires accounting for evolutionary history, genetic diversity, and how likely individuals will adapt to the environmental conditions of the targeted restoration sites (Breed et al., 2019). Here we employ a comprehensive landscape genomic approach to characterize neutral and adaptive genetic variation and provide insights to assist the restoration of a highly degraded mining site and a moderately disturbed Canga highland from the Carajás Mineral Province. We discuss how our results can help define site-adjusted provenancing strategies and argue that our methods can be more broadly applied to assist other restoration and rehabilitation initiatives.

Several studies have stressed the importance of avoiding inbreeding, increasing genetic diversity to maintain evolutionary potential, and minimizing outbreeding depression in restored populations (Broadhurst et al., 2008; Hufbauer et al., 2015; Mijangos et al., 2015; Weeks et al., 2011). The assessment of neutral genetic structure provides information on how to minimize outbreeding depression by avoiding mixing individuals from different evolutionary lineages (Mijangos et al., 2015). Estimates of the genetic neighborhood size, on the other hand, provide clues on how to sample unrelated individuals within seed sourcing areas to increase genetic diversity and reduce the risk of inbreeding depression in restored populations (Breed et al., 2019; Krauss and Koch, 2004). Our initial assessment of neutral genetic structure identified three demographically independent units (or Management Units *sensu* (Funk et al., 2012)), which could be considered distinct provenances to minimize the risk of outbreeding depression (Frankham et al., 2017). Within these zones, our estimates of genetic neighborhood size provide information on within-cluster seed sourcing strategies to maximize genetic diversity. In Serra Sul, for example, seed sources located 5 Km apart are not expected to be related (Fig. S10-S11), and would comprise a better representation of standing genetic variation than individuals collected across smaller spatial scales. Effective population size estimates (Table 1) nevertheless indicate that none of our observed genetic clusters is likely to experience inbreeding depression in the near future based on the 50/500 rule (Jamieson and Allendorf, 2012). Inbreeding levels observed in both species were nonetheless significantly different from zero, suggesting some level of selfing or mating between related individuals is taking place.

Patterns of local adaptation will ultimately determine the ability of plants to effectively colonize and quickly recover disturbed sites (Mijangos et al., 2015). By using the combined candidate loci detected in GEA and GPA, using both univariate and multivariate methods, we improved the detection of single-locus and multi-locus adaptive signals. (Mahony et al., 2019; Talbot et al., 2016; Vangestel et al., 2018). Interestingly, more intersections between GEA and GPA were found when using RDA than when using LFMM (Fig. 3), indicating that most adaptations to local environmental conditions expressing differential phenotypes are polygenic (Forester et al., 2018). Indeed most fitness-related traits in plants have a polygenic basis (Falke et al., 2013), including tolerance to soil with phytotoxic levels of heavy metals (Arnold et al., 2016). We nevertheless note that other genes occurring in the flanking regions of our candidate SNPs could be responsible for the detected adaptive signals, and that many sequences did not match translated proteins, or found matches with uncharacterized proteins. Still, the most frequent amongst our identified candidate proteins are involved in plant defense and stress responses (including reverse transcriptase, ribonuclease H-like domain, P-loop NTPase fold, leucine-rich repeat, and thaumatin) or basic metabolic processes (pentatricopeptide repeat, protein kinase domain, Nitrite/Sulfite reductase ferredoxin-like domain, major intrinsic protein, and kinesin motor domain), suggestive of adaptations to harsh environments (Tables S4 and S5). Some of these putative adaptive genes were shared between the two species as well as with other co-occurring species (Table S5), indicating convergent evolution to similar environmental pressures (Arnold et al., 2016; Yeaman et al., 2016). These shared genes thus constitute primary targets for functional studies investigating the molecular basis of adaptation to Canga environments and minelands.

Cluster-level GEA analyses revealed many cluster-exclusive detections in both species (Fig. S13), suggesting that microgeographic adaptation may play a role in driving genetic patterns within highlands (Richardson et al., 2014). To visualize and better understand the mechanisms behind the observed GEA and GPA we ran additional RDAs using the combined candidate adaptive loci detected in our general LFMM and RDA analyses. As expected, we found similar patterns of GEA in both species (figs. 4a and 4b). Interestingly, the strongest GEA were found with climatic variables in both species, in spite of the coarse resolution of WorldClim data and the narrow climatic variation found across our study area (Fig. S1). Our results thus suggest that local climate constitutes an important environmental filter driving local adaptation, as found in other species from Canga environments (Lanes et al., 2018) and temperate climates (Pais et al., 2017; Pluess et al., 2016). In *M. acutistipula*, Serra Norte populations showed associations with higher SLA, suggesting climatic or soil conditions in Serra Norte are more favorable to plant growth (He et al., 2018). Individuals from Serra da Bocaina and Serra Sul showed associations with lower SLA and lower concentration of several micro- and macro-nutrients, suggesting that increasing leaf thickness in these individuals avoids dissection or better preserves scarce nutrients (Costa-Saura et al., 2016). In contrast, SLA-associations in *D. apurensis* did not separate Canga highlands, showing that the influence of climatic variation on SLA is different across species (Gong and Gao, 2019; Liu et al., 2017). In *D. apurensis*, different genotype associations with leaf micro and macronutrients separated highlands (Fig. 4d), suggesting different physiological requirements or nutrient availability at each site. Low correlations between leaf and soil Fe and Mn concentrations, suggest our study species are controlling nutrient absorption, which makes them suitable for the restoration of areas with a high concentration of these metals. Controlled common-garden or reciprocal transplant experiments are nevertheless needed to assess growth and overall performance of different genotypes (sources) in different soils and climates (Aitken and Bemmels, 2016; Rellstab et al., 2015).

Our local adaptation maps reveal areas containing similar local adaptations (colors) in each species (Fig. 5, Fig. S15), which could be used to delineate seed sourcing strategies. In contrast to the commonly employed Generalized Dissimilarity Models (GDM) (Gugger et al., 2018; Rossetto et al., 2019; Shryock et al., 2015; Supple et al., 2018), our mapping approach based on sPCA allows incorporating GPA and predicting adaptive genetic variation from site-level data, which is particularly useful for areas lacking high-resolution environmental layers. Moreover, sPCA explicitly account for spatial autocorrelation in genetic composition, which is likely to play an important role explaining patterns of local adaptation (Lesica and Allendorf, 1999; Richardson et al., 2014) (see Fig. S16 for alternative adaptation maps generated using GDM). Our adaptation maps showed similar adaptations across Serra da Bocaina (i.e. a single adaptive unit), and most predicted genotypes associated with local environmental samples (climate and soil) clustered together with individuals sampled in Serra da Bocaina. This result indicates that local provenances are probably best adapted to local environmental conditions at this site under contemporary climates (Fig. 6c and 6d), and supports the recommendations made by Lesica & Allendorf (1999) for the restoration of moderately disturbed sites. Since genetic neighborhood size in Serra da Bocaina was roughly 3 Km for both species, our results suggest that local seeds collected in areas separated by at least 3 Km would maximize genetic diversity at this location.

In contrast, predicted genotypes for the environmental data collected at the degraded mine site did not cluster with any of our study populations in either species (Fig. 6a and 6b). This indicates that none of the genotypes we sampled from natural habitats overlap with the multivariate environmental conditions present at the mine site. In this case, mixing genotypes containing different local adaptations could be regarded as the best option to maximize evolutionary potential and facilitate adaptation to novel environments (Lesica and Allendorf, 1999). Seeds could be sourced from all the identified adaptive units (colors in sPCA maps); and within these units they could be sampled in areas separated by the genetic neighborhood size to further enhance genetic diversity. Although mixing individuals from different management units could result in outbreeding depression (Hufford and Mazer, 2003; Weeks et al., 2011), the risk is likely marginal for these study species, which are widely distributed across the continent (Dutra and Morim, 2015; Queiroz, 2015). Moreover, environmental conditions and plant communities show remarkable similarities across the Carajás Mineral Province when compared to other *campo rupestre* formations (Zappi et al., 2019). Such a regional admixture provenancing approach (Bucharova et al., 2019) would represent a significant improvement over introducing exotic species, which are currently being used in mine reclamation programs due to the availability of seeds in large quantities and their ability to quickly colonize mine environments to prevent soil erosion (D’Antonio and Meyerson, 2002; Gastauer et al., 2019; Silva et al., 2018). Our results could guide the establishment of seed production areas for both native species, aiming to overcome shortfalls in seed availability while capturing standing neutral and adaptive genetic variation (Nevill et al., 2016).

Our work illustrates how neutral and adaptive genetic variation can be used to provide evidence-based recommendations for provenance schemes aimed to effectively restore sites ranging between moderately disturbed and highly degraded. In our two study species, local provenances were found optimal to restore a moderately disturbed site, whereas a mixture of genotypes was suggested as the most promising strategy to recover a highly degraded mining site, to which local provenances were not adapted. Our proposed methodological approach (Fig. 1) can be more broadly applied to define site-adjusted provenance strategies in other locations and for other disturbance regimes. We recognize that the high costs associated with genomic analyses and the complexity of bioinformatic and statistical analyses represent important barriers for practitioners (Breed et al., 2019; Shafer et al., 2015). Still, as genomic data becomes available for more species exhibiting different life-history characteristics, restoration genomic initiatives using similar methods coupled with visually appealing and user-friendly interfaces (Rossetto et al., 2019), are likely to substantially improve restoration outcomes.

## Acknowledgments

Funding was provided by Instituto Tecnológico Vale, Conselho Nacional de Desenvolvimento Científico e Tecnológico (CNPq) grants 301616/2017-5 (RJ), 153535/2018-0 (MG) and 316067/2018-0 (LCRM), and Coordenação de Aperfeiçoamento de Pessoal de Nível Superior (CAPES) grant 88887.156652/2017-00 (CSC). We thank Cesar Neto and Eder Lanes for assistance in the field, Nelson Carvalho and Santelmo Vasconcelos for assistance in genome size estimation, Eder Lanes and Manoel Lopes for help in the laboratory, and three anonymous referees for improving earlier versions of this manuscript.

## Data Availability Statement

Geographic coordinates, genotypes in Variant Call Format, and sequences in FASTA format for both species are available in Figshare: https://doi.org/10.6084/m9.figshare.12185235.v1. All the mentioned R scripts have been deposited in GitHub and their url addresses provided in the text.

## Author contributions

RJ conceived, designed and coordinated the project. CSC, MG and RJ coordinated the field work and sampling. CSC, SM and MG performed laboratory work. CSC, LCRM, LT, BF, MG and RJ performed the data analysis. The first draft of the paper was written by CSC and RJ and all authors contributed to discussing the results and editing the paper.

