## Supporting Information for "Combining genotype, phenotype, and environmental data to delineate site-adjusted provenance strategies for ecological restoration"

The following Supporting Information is available for this article:

**Methods S1:** Additional methods on environmental and phenotype data, genome size estimation, genotype-by-sequencing and bioinformatic processing.
**Fig. S1:** Maps of the bioclimatic layers used in GEA analyses.
**Fig. S2:** Correlations between environmental and phenotype predictor variables.
**Fig. S3-S8:** Histograms of test significance values (*p*-values) for GEA and GPA analyses, using the calculated and modified genomic inflation factor (GIF).
**Fig. S9:** Plots showing the optimal number of genetic clusters (k) for each species, based on the *snmf* function from the LEA package and DAPC from adegenet .
**Fig. S10-S11:** Fine-scale spatial genetic structure for *Mimosa acutistipula* var. *ferrea* and *Dioclea apurensis*.
**Fig. S12:** Global Redundancy analysis (RDA) to assess genotype-environment associations and genotype-phenotype associations in each species.
**Fig. S13:** Venn diagrams showing the intersection of contigs deteced in general GEA analyses and in cluster-level GEA analyses for each species.
**Fig. S14:** Barplot showing eigenvalues of sPCA on all candidate loci found in each species.
**Fig. S15:** Spatial distribution of adaptive genetic variation in each species, based on sPCA on intersected candidate loci.
**Fig. S16:** Spatial distribution of adaptive genetic variation in each species, based on Generalized Dissimilarity Models (GDM).
**Table S1:** Eigenvalues and percentage of total variance explained by each PCA axis retained to select environmental and phenotypic predictor variables in each species.
**Table S2:** Summary of the number of adaptive signals detected using genotype-environment association (GEA) in each species.
**Table S3:** Summary of the number of adaptive signals detected using genotype-phenotype association (GPA) in each species.
**Table S4:** Annotated candidate proteins detected using GEA and GPA in each species.
**Tables S5:** Annotated candidate proteins shared among different plant species occurring in our study region.

**Methods S1**

*Environmental data*

Chemical analyses of soil samples included pH (measured in $H_2O$, 1:2.5 mass:volume), organic matter (MO, determined by colorimetric method using $Na_2Cr_2O_7$ and $H_2SO_4$ extractors), available P, K, and Na (extracted by the Mehlich 1 chemical method, 0.05 mol $L^{-1}$ HCl, 0.0125 mol $L^{-1}$ $H_2SO_4$ and detected by colorimetry [P], flame photometry [K] or Inductively Coupled Plasma-Atomic Emission Spectometry [ICP-AES, Na]), exchangeable Ca, Mg, and Al (extracted with KCl 1 M and determined by Atomic Absorption Spectrometry), available B (determined by spectrophotometry using heating and $BaCl_2$ 1.25 g $L^{-1}$), exchangeable S (measured by turbidimetry using $Ca(H_2PO_4)2.H_2O$ 0.01 M) and available Cu, Fe, Mn and Zn (determined using diethylene triamine pentaacetic acid method) (Silva, 2009).

*Phenotypic data*

Leaf tissue samples were oven-dried at 60ºC until they maintained a constant weight and were processed in a Willey mill to obtain a fine powder. Inductively Coupled Plasma Atomic Emission Spectroscopy (ICP-AES) was then employed to determine leaf nutrient content, relying on sulfuric digestion to quantify nitrogen, nitro-perchloric digestion to quantify other nutrients, and a dry digestion to quantify boron. To estimate the specific leaf area (SLA, in $cm^2$ $g^{-1}$), leaflets were scanned using a V750 PRO Epson scanner, analyzed using the ImageJ software (Schneider *et al.*, 2012), oven-dried (60ºC) until reach constant weigh, and weighed on a precision balance (± 0.0001 g) (Pérez-Harguindeguy *et al.*, 2013).

*Genome size estimation*

We used flow cytometry to estimate genome size in both species. Nuclei were obtained from fresh leaf tissues chopped along with references in general purpose buffer with 1% Triton X-100 and 1% PVP-30 (Loureiro *et al.*, 2007). The whole sample preparation was conducted on ice until the events acquisition on a PI fluorescence mean under a 575/26 bandpass filter. Triplicates of 1000 PI stained nuclei were analyzed under a 488 nm laser on BD FACS Aria II cytometer. The internal standard used was tomato (*Lycopersicon esculentum*; 2C = 1.98 pg).

**Figures**

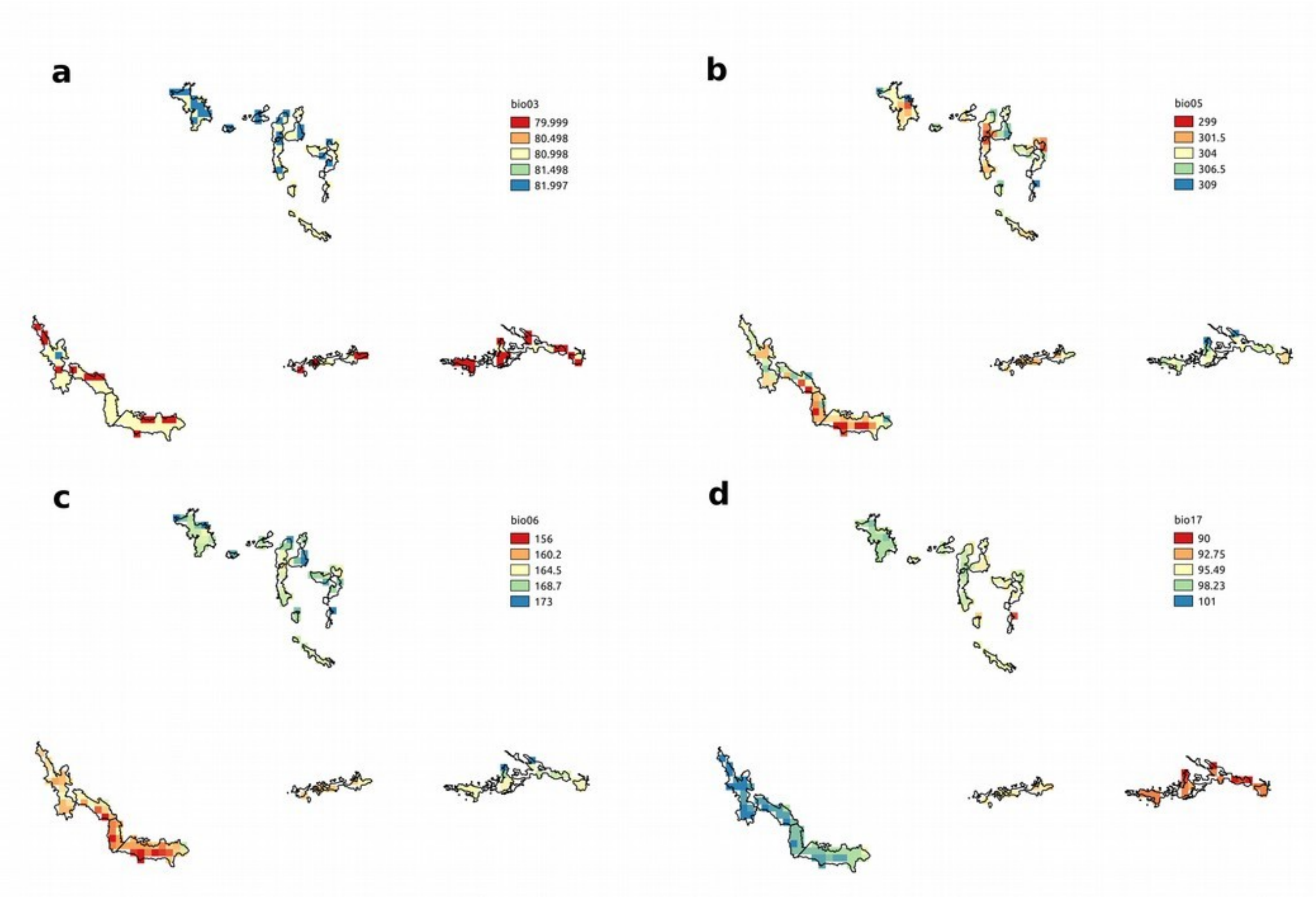

**Figure S1:** Maps of the bioclimatic layers used in GEA analyses. These were isothermality (bio03, **a**), maximum temperature of the warmest month (bio05, **b**), minimum temperature of the coldest month (bio06, **c**), and precipitation of the driest quarter (bio17, **d**).

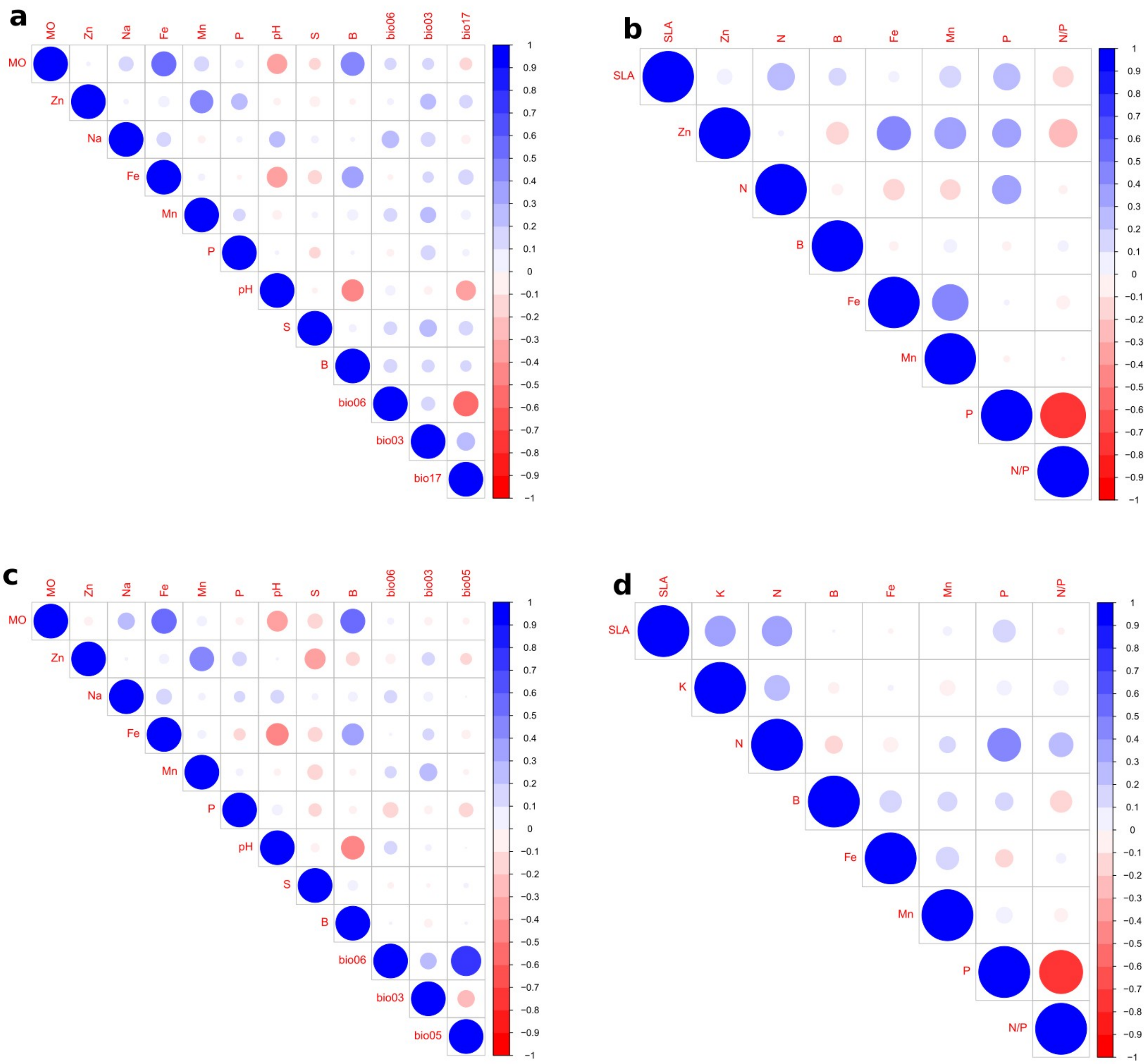

**Figure S2**: Correlations between environmental predictor variables (left panels- MO, Zn, Na, Fe, Mn, P, pH, S, B, bio03, bio05, bio06, bio17) and phenotype predictor variables (right panels- SLA, K, Zn, N, B, Fe, Mn, P, N/P) for *Mimosa acutistpula* var. *ferrea* (a and c) and *Dioclea apurensis* and (b and d). Colors indicate the direction (red negative and clue positive) and size the magnitude of Pearson's correlation coefficients.

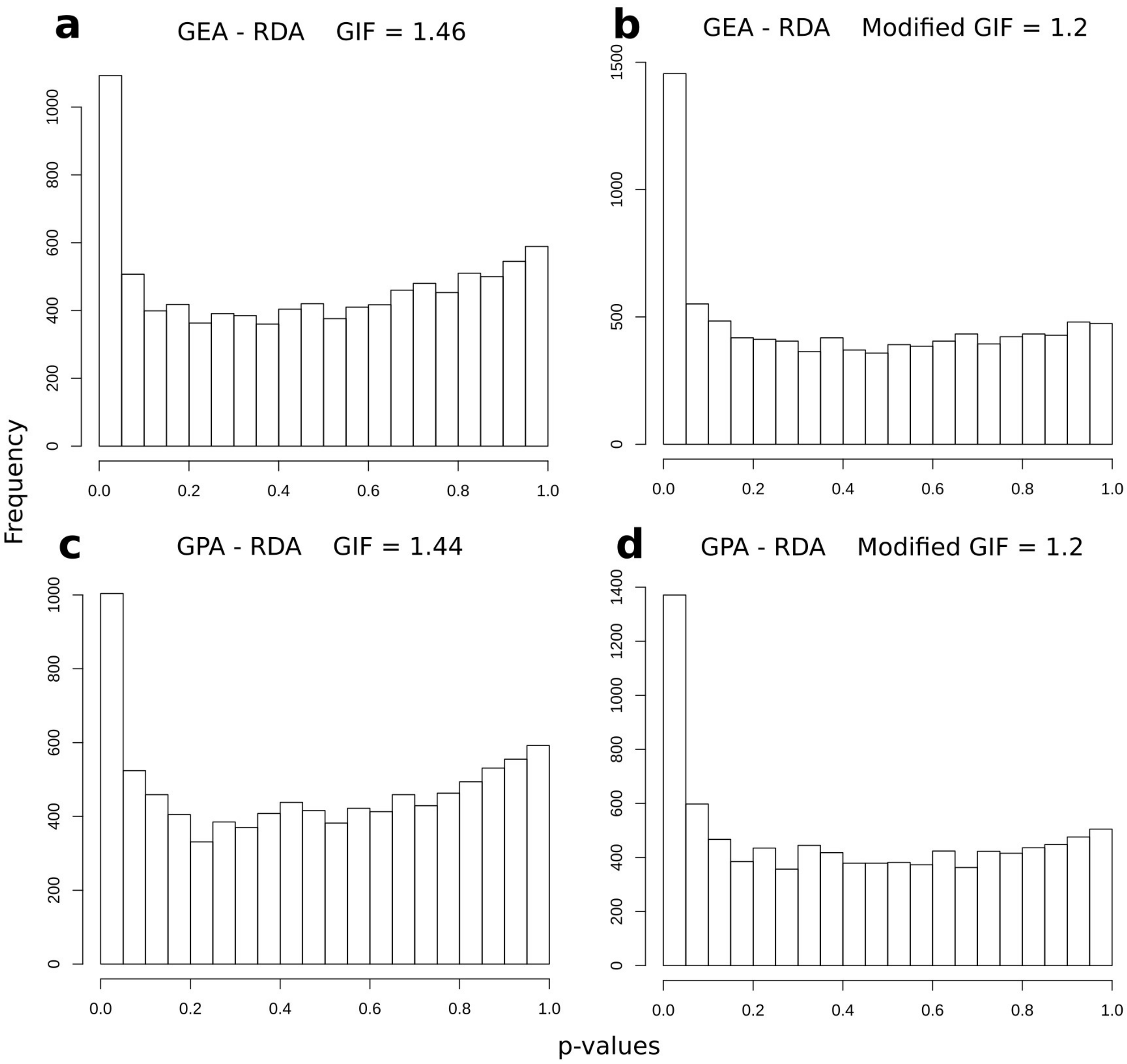

**Figure S3:** Histograms of test significance values (*p*-values) for GEA-RDA (upper panels) and GPA-RDA analyses (lower panels) in *Mimosa acutistipula* var. *ferrea,* using the calculated (**a**, **c**) and modified (**b**, **d**) genomic inflation factor (GIF).

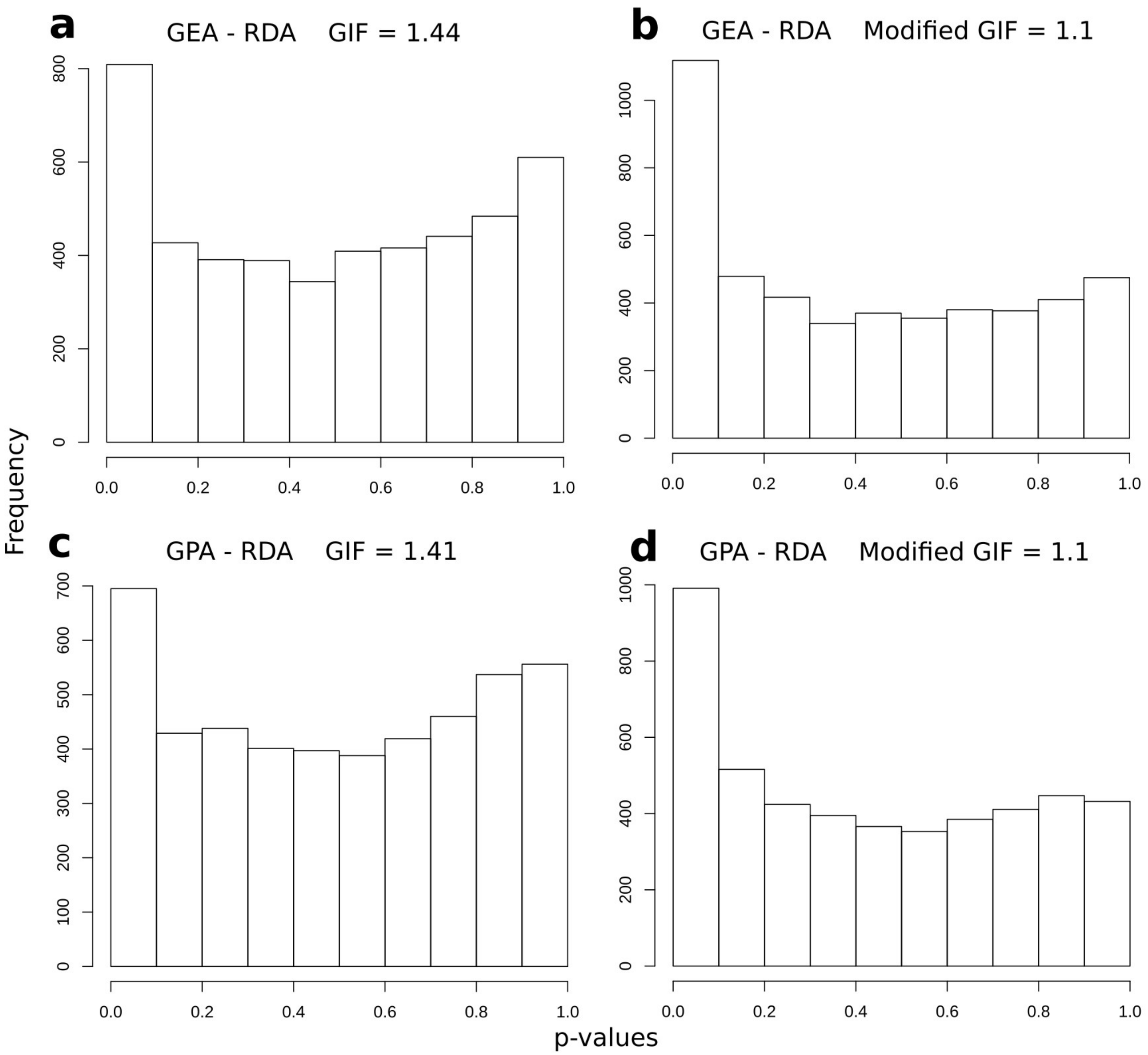

**Figure S4:** Histograms of test significance values (*p*-values) for GEA-RDA (upper panels) and GPA-RDA analyses (lower panels) in *Dioclea apurensis,* using the calculated (**a**, **c**) and modified (**b**, **d**) genomic inflation factor (GIF).

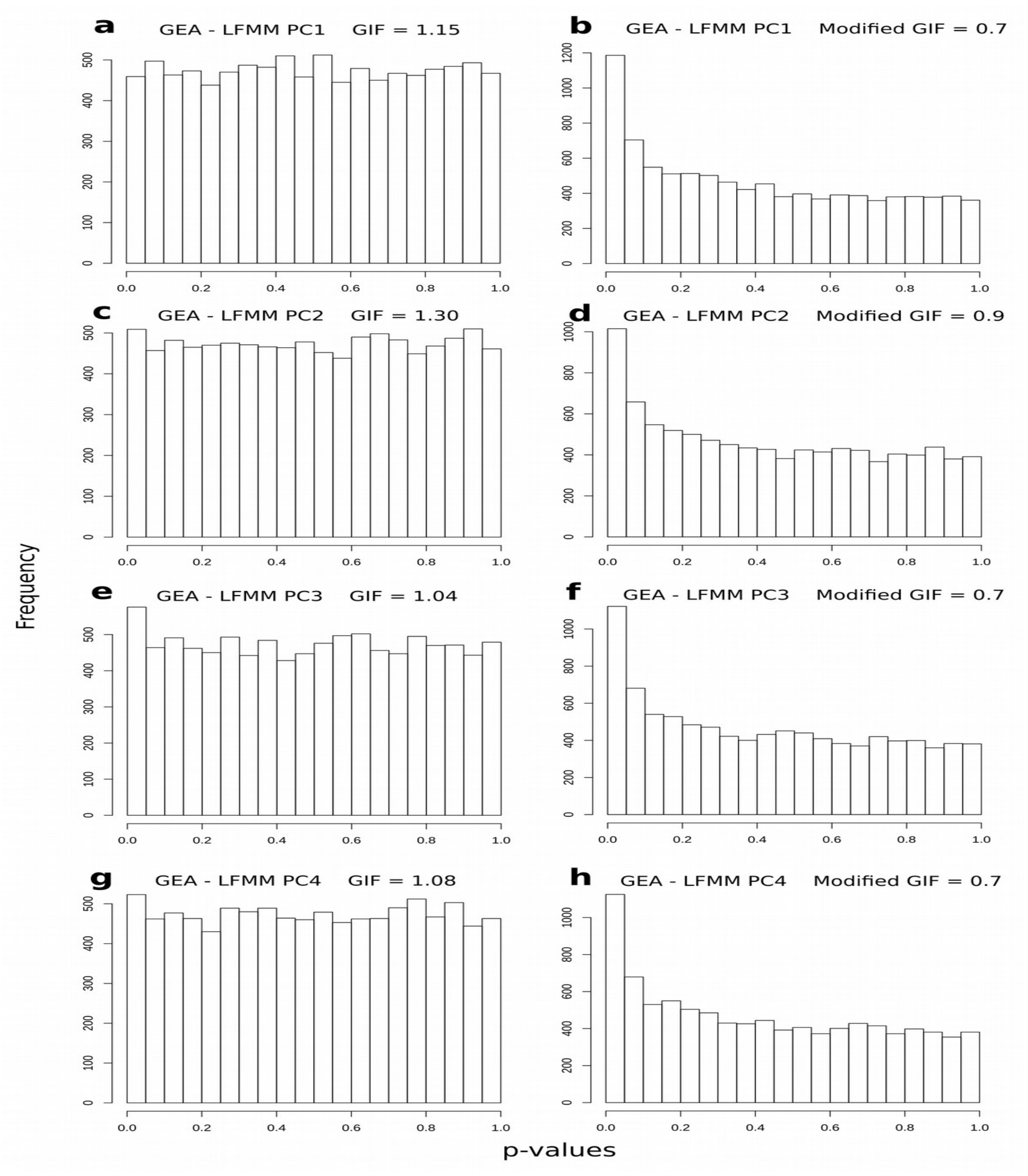

**Figure S5:** Histograms of test significance values (*p*-values) for GEA-LFMM analyses in *Mimosa acutistipula* var. *ferrea*, using the calculated (**a**, **c, e, g**) and modified (**b**, **d, f, h**) genomic inflation factor (GIF). Each line represents a principal component.

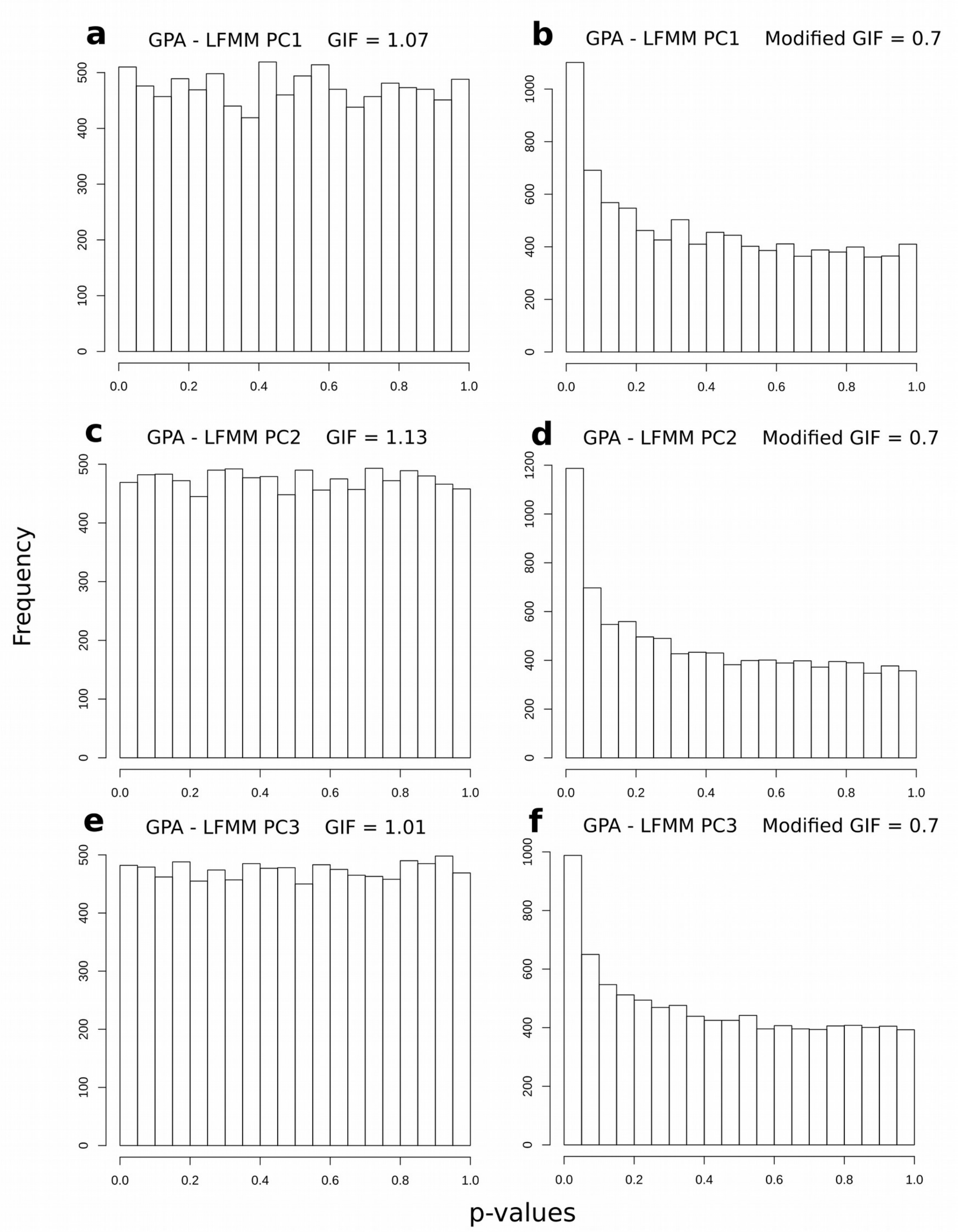

**Figure S6:** Histograms of test significance values (*p*-values) for GPA-LFMM analyses in *Mimosa acutistipula* var. *ferrea*, using the calculated (**a**, **c, e**) and modified (**b**, **d, f**) genomic inflation factor (GIF). Each line represents a principal component.

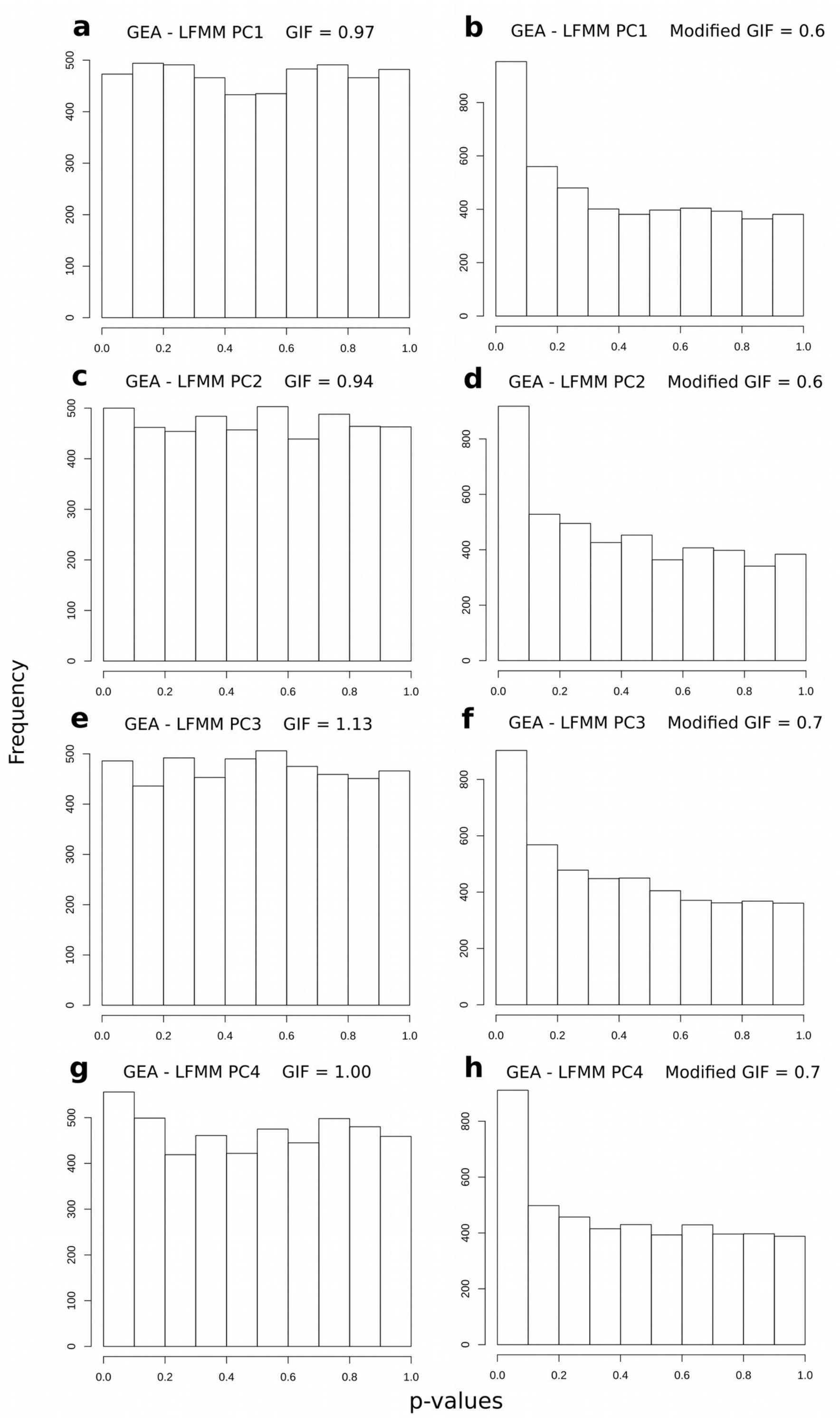

**Figure S7:** Histograms of test significance values (*p*-values) for GEA-LFMM analyses in *Dioclea apurensis*, using the calculated (**a**, **c, e, g**) and modified (**b**, **d, f, h**) genomic inflation factor (GIF). Each line represents a principal component.

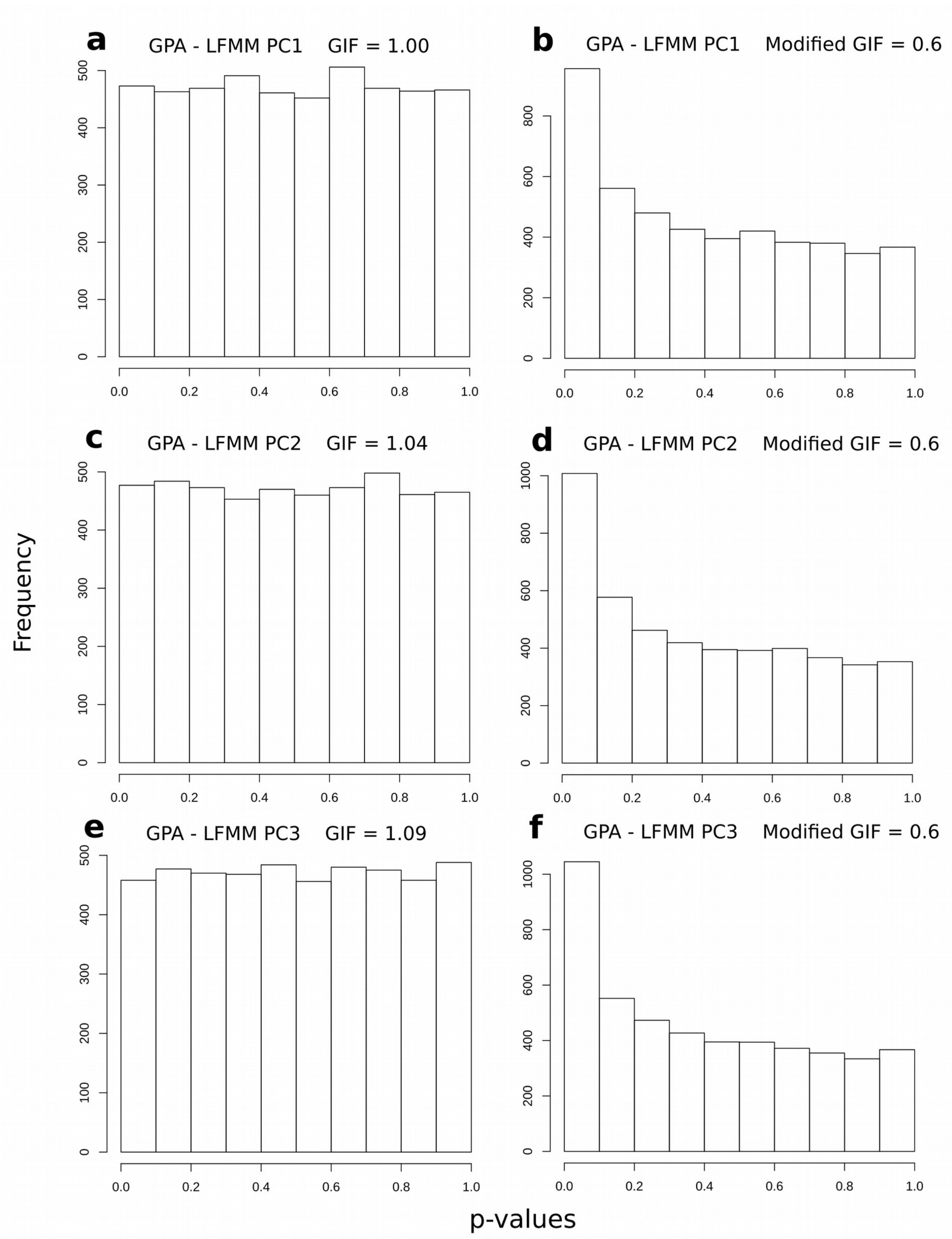

**Figure S8:** Histograms of test significance values (*p*-values) for GPA-LFMM analyses in *Dioclea apurensis*, using the calculated (**a**, **c, e**) and modified (**b**, **d, f**) genomic inflation factor (GIF). Each line represents a principal component.

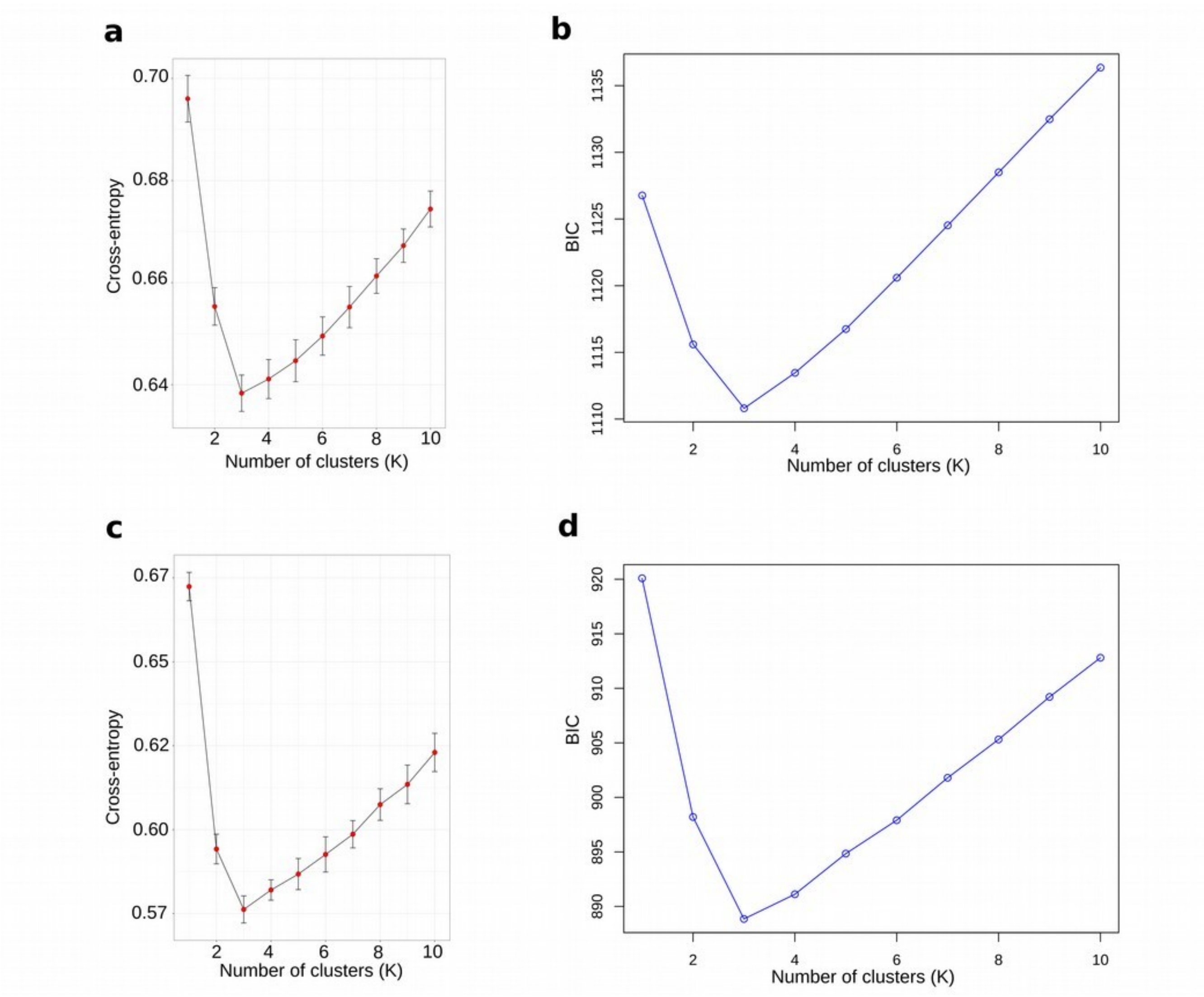

**Figure S9:** Plots showing the optimal number of genetic clusters (k) for *Mimosa acutistipula* var. *ferrea* (a, b) and *Dioclea apurensis* (c, d) based on the *snmf* function from the LEA package (left panels) and DAPC from adegenet (right panels).

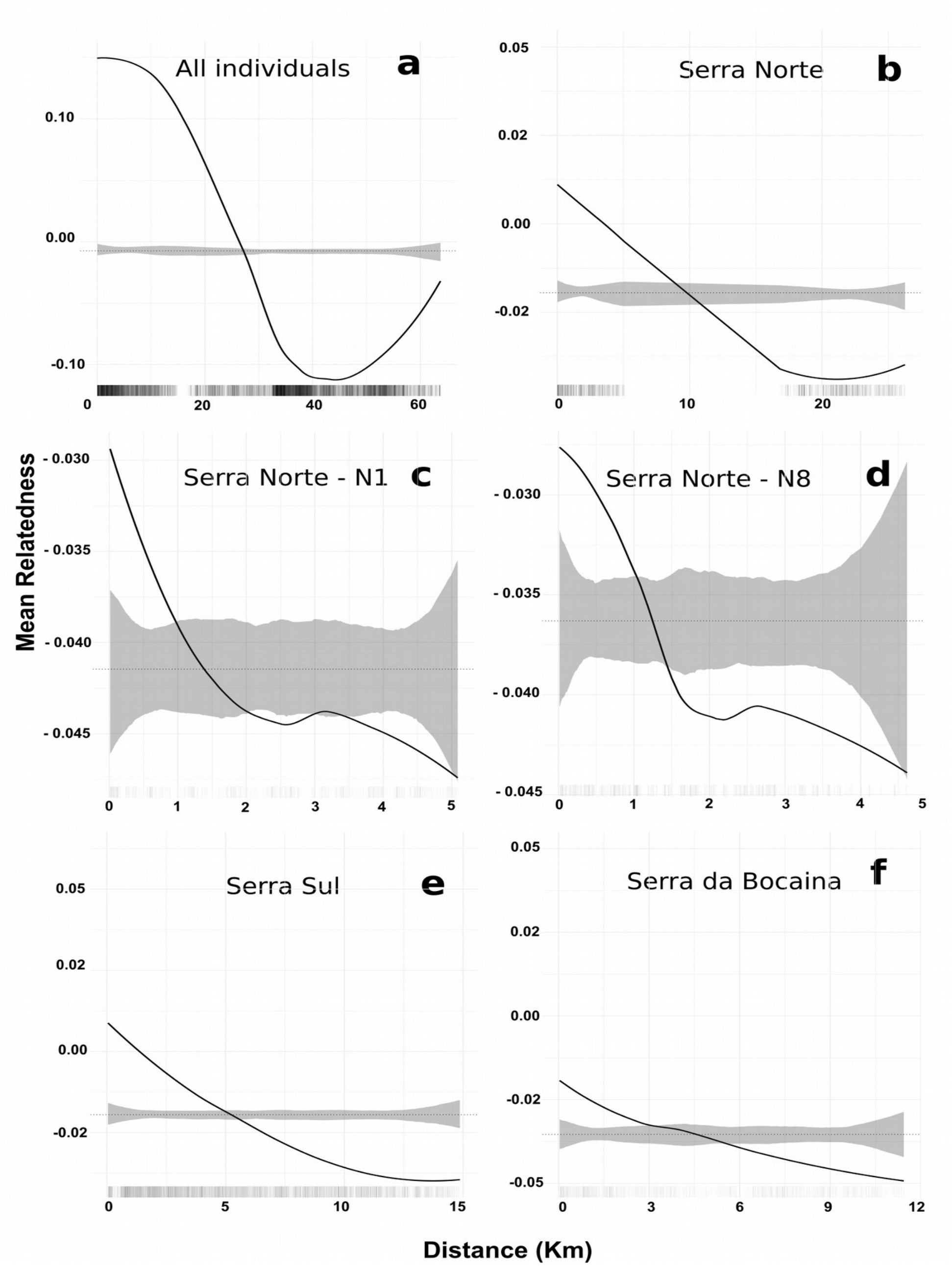

**Figure S10:** Fine-scale spatial genetic structure for *Mimosa acutistipula* var. *ferrea* using all individuals (a), and using individuals from each genetic cluster (b, e and f). Due to sampling discontinuities we also assessed fine-scale spatial genetic structure separately in Serra Norte (c, and d). Black solid lines are the LOESS fit to the observed relatedness, and gray shaded regions are 95% confidence bounds around the null expectation (black dotted lines). The genetic neighborhood size is represented by the distance at which genetic relatedness stops being spatially autocorrelated (where black lines cross shaded regions).

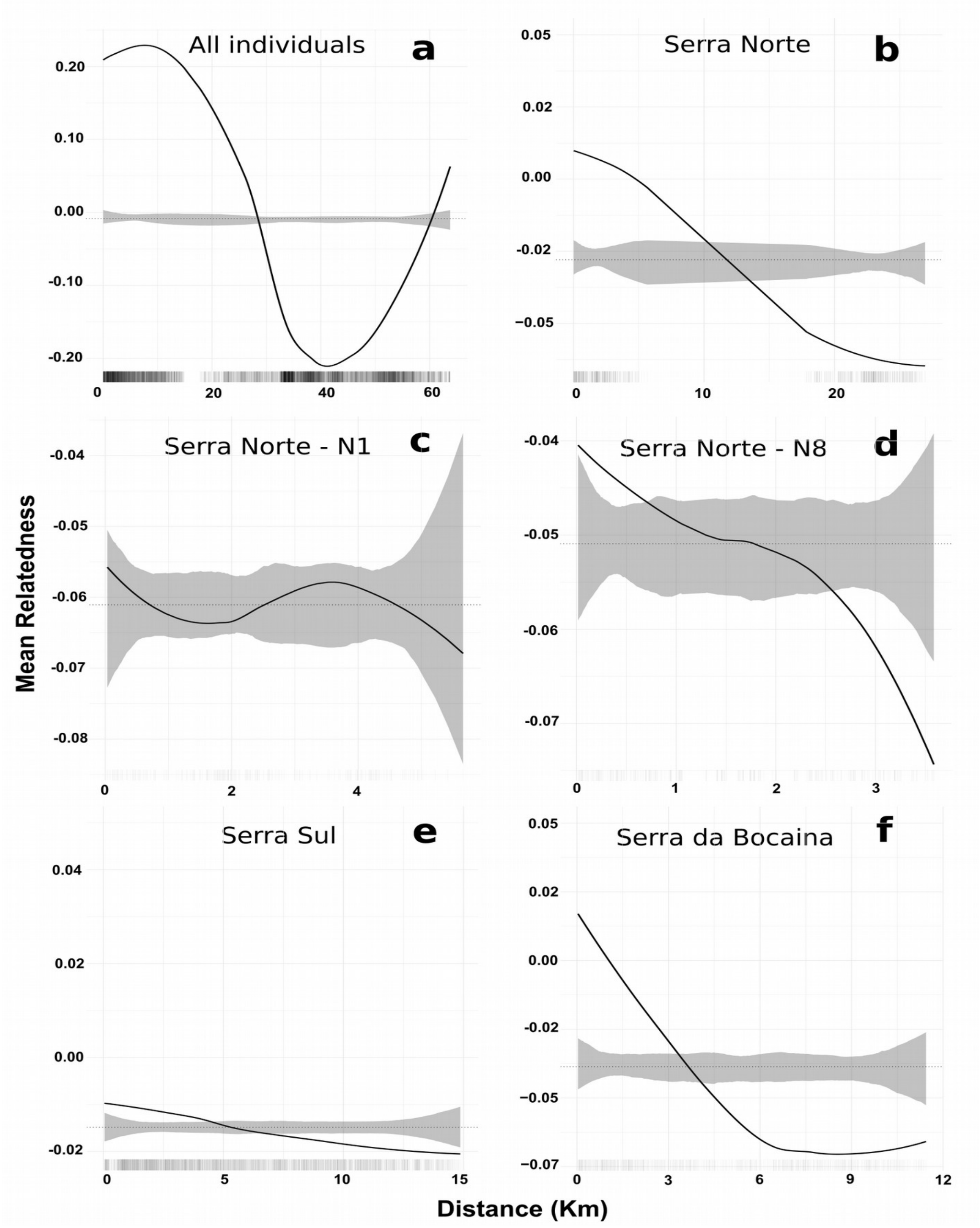

**Figure S11:** Fine-scale spatial genetic structure for *Dioclea apurensis* using all individuals (a), and using individuals from each genetic cluster (b, e and f). Due to sampling discontinuities we also assessed fine-scale spatial genetic structure separately in Serra Norte (c, and d). Black solid lines are the LOESS fit to the observed relatedness, and gray shaded regions are 95% confidence bounds around the null expectation (black dotted lines). The genetic neighborhood size is represented by the distance at which genetic relatedness stops being spatially autocorrelated (where black lines cross shaded regions).

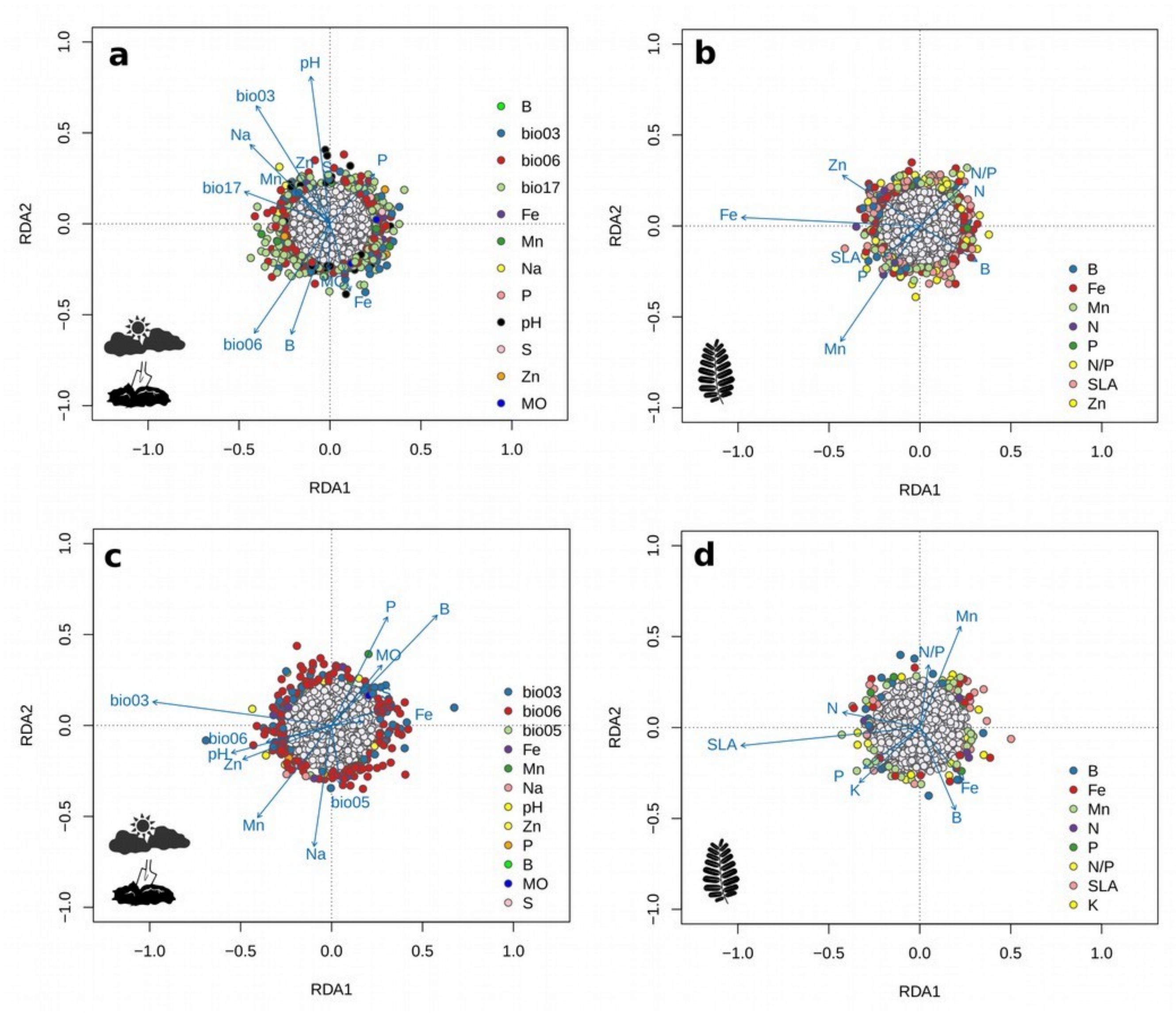

**Figure S12:** Global Redundancy analysis (RDA) for *Mimosa acutistipula* var. *ferrea* (**a** and **b**) and *Dioclea apurensis* (**c** and **d**) to assess genotype-environment associations (**a** and **c**) and genotype-phenotype associations (**b** and **d**). Candidate loci were selected based on Mahalanobis distance D on loci loadings from the RDA analysis, and colors represent the variables showing the strongest correlations with candidate SNPs. Neutral (not candidate) SNPs are shown in grey, and blue arrows represent the environmental and phenotypic predictors.

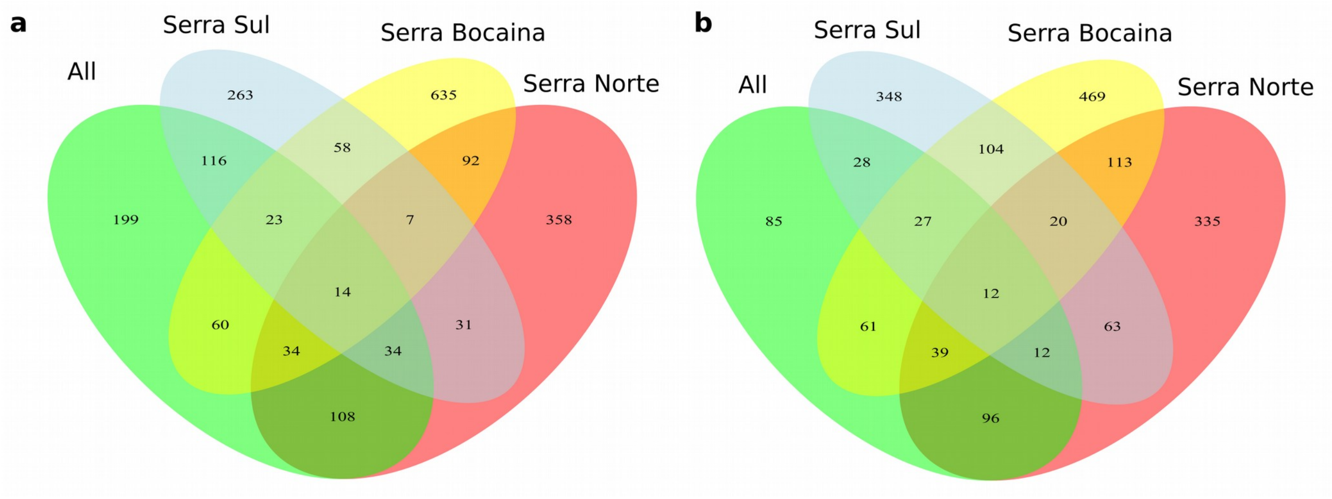

**Figure S13:** Venn diagram showing the intersection of contigs (RAD-tags) containing candidate SNPs for *Mimosa acutistipula* var. *ferrea* (a) and *Dioclea apurensis* (b) deteced in general GEA analyses (using all individuals) and in cluster-level GEA analyses (using only individuals belonging to the same cluster, see methods for details).

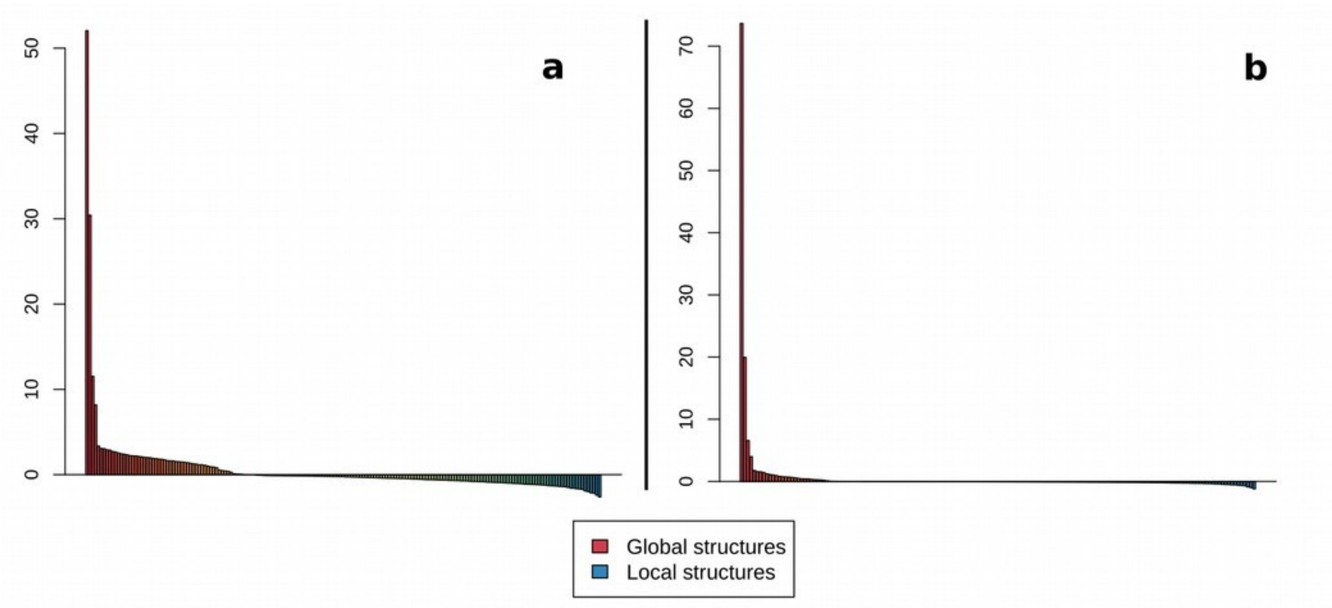

**Figure S14:** Barplot showing eigenvalues of sPCA on all candidate loci found in *Mimosa acutistipula* var. *ferrea* (**a**) and *Dioclea apurensis* (**b**). While global structure is represented as positive spatial auto-correlation, local structure is shown with negative values.

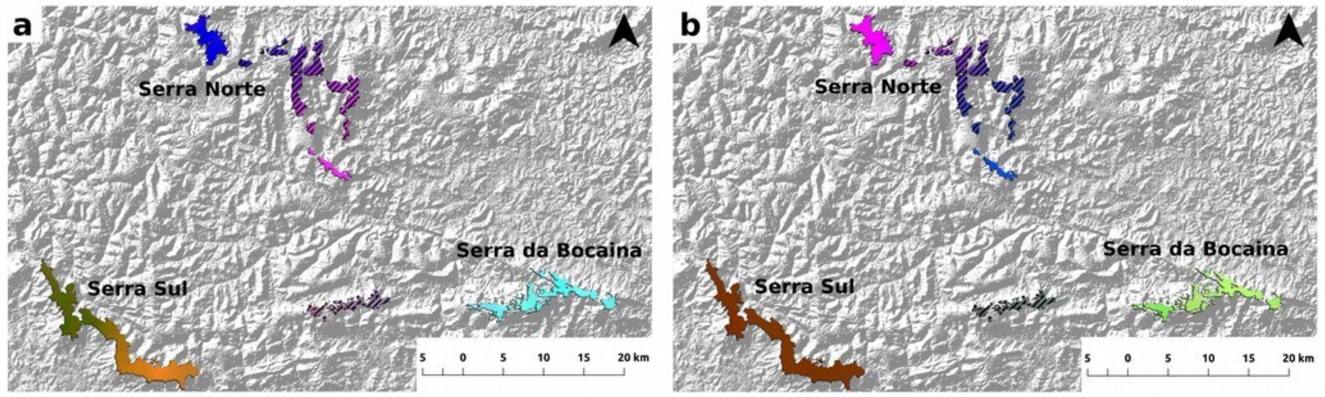

**Figure S15:** Spatial distribution of adaptive genetic variation in *Mimosa acutistipula* var. *ferrea* (**a**) and *Dioclea apurensis* (**b**) based on sPCA on intersected candidate loci. The maps represent an RGB composite made using interpolated principal components from a sPCA, ran on the intersected loci (found in both GEA and GPA). Regions with similar colors within each panel represent analogous genetic composition and areas with diagonal lines were not sampled (i.e., adaptive genetic composition was extrapolated from neighboring areas).

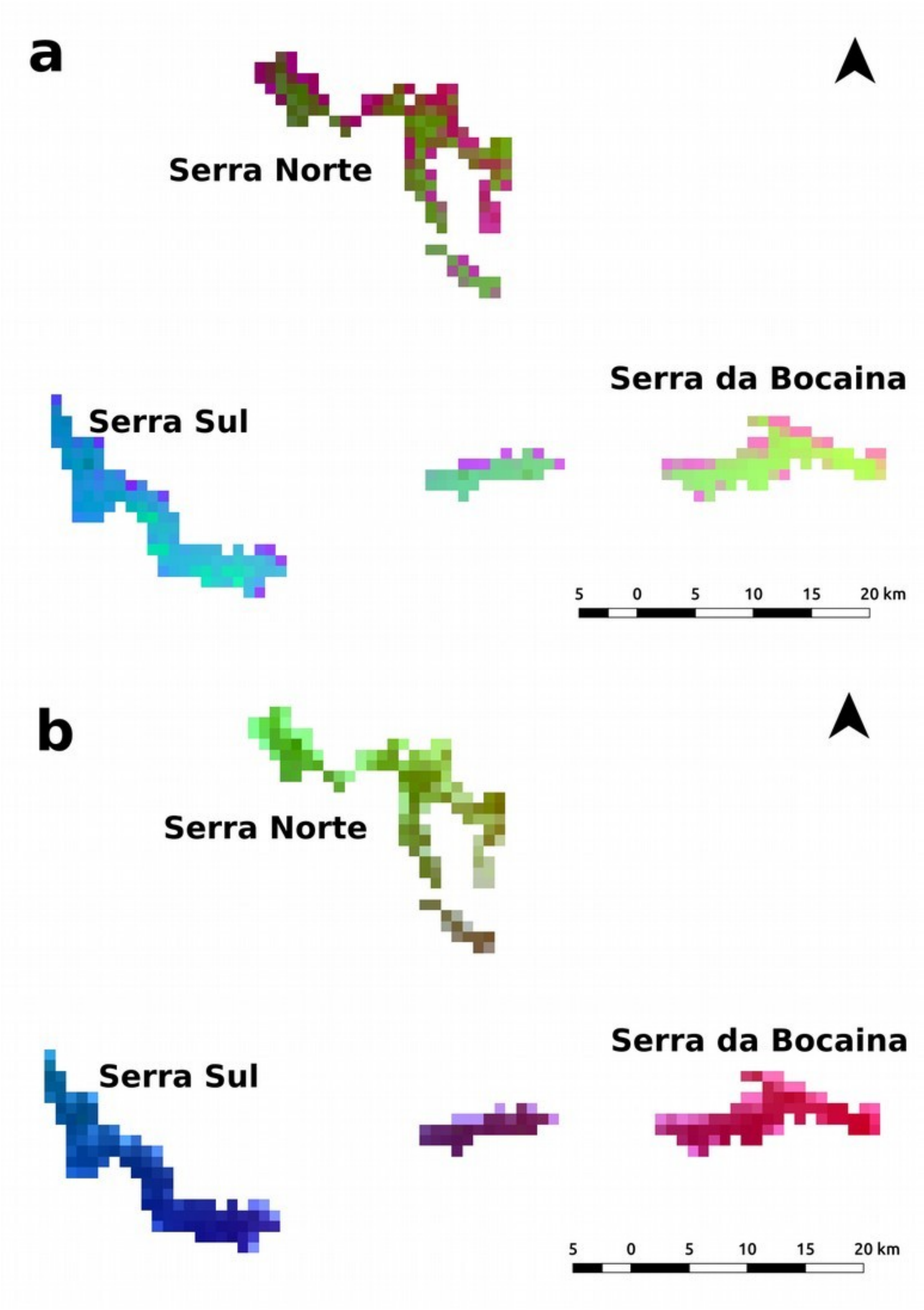

**Figure S16:** Spatial distribution of adaptive genetic variation in *Mimosa acutistipula* var. *ferrea* (**a**) and *Dioclea apurensis* (**b**) based on Generalized Dissimilarity Models (GDM). Regions with similar colors represent analogous genetic compositions. When compared with sPCA (Figs. 5 and S15) GDM seem to overemphasize the effect of environmental variables and overestimate genetic dissimilarity (spatial autocorrelation is expected to dissolve stark boundaries).

**Tables**

**Table S1:** Eigenvalues and percentage of total variance explained by each PCA axis retained to select environmental and phenotypic predictor variables in *Mimosa acustistipula* and *Dioclea apurensis*.

| **Species** | **Variable type** | **Axis** | **Eigenvalue** | **Variance (%)** |
|---|---|---|---|---|
| *M. acutistipula* | Soil | 1 | 3.8 | 26 |
| | | 2 | 2.2 | 15 |
| | | 3 | 1.4 | 10 |
| | Climatic | 1 | 10.0 | 53 |
| | | 2 | 3.26 | 17 |
| | | 3 | 2.2 | 12 |
| | Phenotype | 1 | 3.0 | 28 |
| | | 2 | 1.8 | 17 |
| | | 3 | 1.1 | 10 |
| *D. apurensis* | Soil | 1 | 3.8 | 26 |
| | | 2 | 2.4 | 17 |
| | | 3 | 1.5 | 10 |
| | Climatic | 1 | 10.3 | 55 |
| | | 2 | 3.2 | 17 |
| | | 3 | 2.2 | 12 |
| | Phenotype | 1 | 2.5 | 23 |
| | | 2 | 1.6 | 15 |
| | | 3 | 1.3 | 12 |

**Table S2:** Summary of the number of adaptive signals detected in genotype-environment association (GEA) analyses for *Mimosa acutistipula* var. *ferrea* and *Dioclea apurensis*.

| Species | Signal Type | Total analyzed | Total candidates LFMM | Total candidates RDA | Shared (LFMM-RDA) | LFMM | | | | RDA | | | | | | | | | | | | |
|---|---|---|---|---|---|---|---|---|---|---|---|---|---|---|---|---|---|---|---|---|---|---|
| | | | | | | PC1 | PC2 | PC3 | PC4 | OM | Zn | Na | Fe | Mn | P | pH | S | B | Bio06 | Bio03 | Bio17 | Bio05 |
| *M. acutistipula* | SNPs | 9480 | 208 | 430 | 7 | 70 | 40 | 55 | 55 | 2 | 5 | 4 | 5 | 9 | 2 | 31 | 7 | 5 | 115 | 54 | 191 | - |
| | Contigs | 5678 | 198 | 403 | 13 | - | - | - | - | - | - | - | - | - | - | - | - | - | - | - | - | - |
| *D. apurensis* | SNPs | 4720 | 161 | 231 | 17 | 44 | 57 | 35 | 32 | 1 | 5 | 2 | 6 | 2 | 2 | 4 | 1 | 1 | 156 | 48 | - | 3 |
| | Contigs | 3551 | 154 | 225 | 19 | - | - | - | - | - | - | - | - | - | - | - | - | - | - | - | - | - |

**Table S3:** Summary of the number of adaptive signals detected in genotype-phenotype association (GPA) for *Mimosa acutistipula* var. *ferrea* and *Dioclea apurensis*.

| Species | Signal Type | Total analyzed | Total candidates LFMM | Total candidates RDA | Shared (LFMM-RDA) | LFMM | | | RDA | | | | | | | | |
|---|---|---|---|---|---|---|---|---|---|---|---|---|---|---|---|---|---|
| | | | | | | PC1 | PC2 | PC3 | SLA | K | Zn | N | B | Fe | Mn | N/P | P |
| *M. acutistipula* | SNPs | 9480 | 96 | 295 | 6 | 43 | 43 | 10 | 83 | 0 | 11 | 17 | 21 | 67 | 54 | 37 | 5 |
| | Contigs | 5678 | 94 | 281 | 7 | - | - | - | - | - | - | - | - | - | - | - | - |
| *D. apurensis* | SNPs | 4720 | 190 | 125 | 15 | 58 | 75 | 61 | 26 | 13 | 0 | 6 | 18 | 18 | 33 | 4 | 7 |
| | Contigs | 3551 | 185 | 119 | 16 | - | - | - | - | - | - | - | - | - | - | - | - |
